# Quantitative Estimation of Long-Range Interactions at the Nanoscale

**DOI:** 10.1101/478800

**Authors:** Vishesh Sood, Sunandan Dhar, Dhirendra S. Katti

## Abstract

Nano-bio interfaces attune nanoparticle-mediated biological responses. The nano-bio interface, like all interfacial interactions, is governed by non-covalent long-range interactions (LRIs). These LRIs include electrostatic, electrodynamic and acid-base interactions. There is a lack of understanding about the contribution of LRIs at the nano-bio interface for want of suitable methods for the estimation of dispersive, acidic, and basic components of the surface tension of nanoparticles. To address this, we developed an experimental and theoretical framework for the estimation of surface tension components of nanoparticles and biomacromolecules by partitioning them in a biphasic system. The work presented here is the first instance in the literature for estimating the surface tension components of nanoparticles and biomacromolecules suspended in aqueous suspensions. We also observed that LRIs have a deterministic role in biologically relevant phenomena such as salt-induced nanoparticle agglomeration and protein-nanoparticle interaction. Collectively, the results presented in this work provide a rapid and inexpensive framework for predicting the energetics of nanoparticle-nanoparticle and nanoparticle-protein interactions by estimating average ensemble surface characteristics like surface tension and surface charge density.

---

Nanoparticles are ubiquitous in nature due to a plethora of natural and anthropogenic sources. Nano-bio interfaces are established by the propensity of nanoparticles to interact with biological systems present in their vicinity owing to their high surface energy^1^. Nano-bio interactions play a significant role in nanoparticle-mediated biological responses including adsorption of specific proteins^2^ and consequential nanoparticle uptake^3, 4^, perturbation of cellular homeostasis^5^, inflammation^6^, or controlled biodistribution^7^. However, there is an inadequate understanding of the fundamental aspects that govern the interaction between nanoparticles and biological systems.

The non-covalent long-range interactions (LRIs) primarily govern the interfacial interactions between materials and biological systems such as microorganisms and bacteria^8, 9^. The interfacial LRIs include electrostatic (EL) or Coulombic interactions, electrodynamic (van der Waal’s forces) or Lifshitz-van der Waals (LW) interactions, and polar (structural surface forces) or acid-base (AB) interactions ^10, 11^. EL, LW, and AB interactions together constitute primary LRIs at nanoscale^12^. The quantification of LRIs has been shown to aid in predicting protein adsorption on micron-sized silica particles and hence provide meaningful information about the implication of material-bio interfaces on biological functions^13^. However, in the context of nanoparticles, the role of only EL interactions is understood as the surface charge density of nanoparticles can readily be estimated by measuring their zeta potential^14, 15^. Whereas, the literature is scarce on the contribution of LW and AB interactions between nanomaterials and biomacromolecules primarily due to non-availability of methods. To address this, a biological surface adsorption index based approach was developed which provided an insight into the fundamental nature of interactions governing nanoparticle-biomacromolecule interactions^16, 17^. The proposed index, however, provided only relative information, and the index had limited utility for the quantification of the contribution of LRIs. Contact angle measurement is the method of choice for the quantification of LW and AB interfacial energies between planar surfaces^18^. Macroscopic contact angle measurement, however, is not suitable for nanoparticles as sample preparation results in micron-scale heterogeneity of the surface^19^. To circumvent this, significant effort has gone into measuring microscopic contact angle using different techniques^20, 21, 22, 23, 24^. Most of these methods provide limited information as they either provide a single contact angle, require extensive sample preparation or expensive equipment. Another alternative is to use biphasic partitioning to predict the biological behavior of molecules. For example, octanol-water partitioning provides a straightforward estimate of the lipophilicity of a compound^25^. The lipophilicity of a compound plays a deterministic role in its ability to cross the blood-brain barrier and thus could predict its biodistribution inside the body^26^. However, partitioning coefficient of nanoparticles does not provide any significant information about the biological behavior of the nanoparticles^27^.

To elucidate meaningful information from biphasic partitioning data of nanoparticles or biomacromolecules, we developed a theoretical and experimental framework correlating the biphasic partitioning of nanoparticles or biomacromolecules to their surface tension components (**Figure 1**). Further, we used the estimated surface tension components to understand the contributions of LRIs including EL, LW and AB interactions in biologically relevant processes like salt-induced nanoparticle agglomeration and nanoparticle-protein interactions (**Figure 1**).

**Figure 1.**
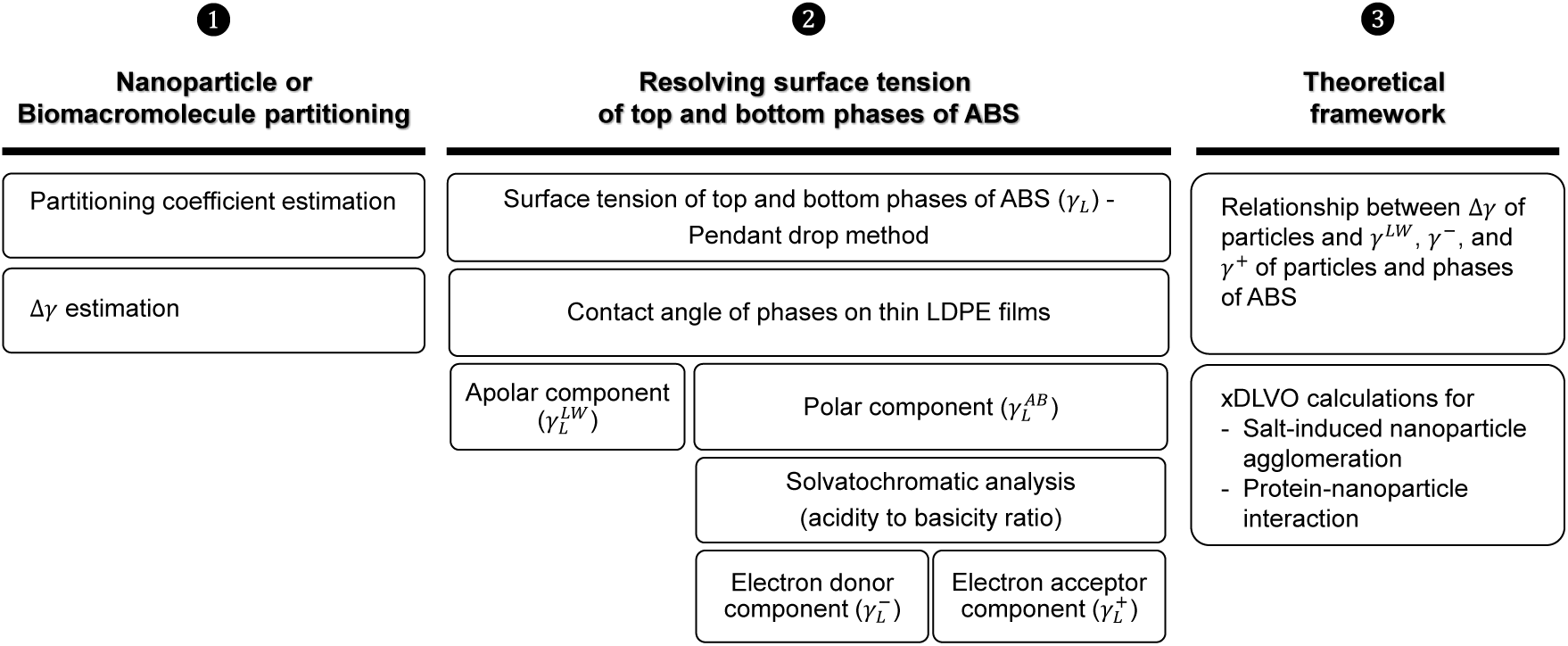
Outline of the methodology used for quantitative estimation of non-covalent long-range interactions (LRIs) at nanoscale. In Step ❶, nanoparticles or biomacromolecules were partitioned in aqueous biphasic systems (ABS) to calculate partitioning coefficient and interfacial interaction parameter (∆*γ*). In Step ❷, the surface tension of polymeric phases was resolved into dispersive 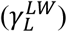 and polar 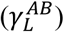 components using contact angle on an apolar surface (thin LDPE film). The polar component was resolved further into electron donor 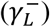 and acceptor 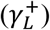 components using solvatochromic analysis. In step ❸, a theoretical framework was developed for the relationship between ∆*γ* of particles (nanoparticles or biomacromolecules) estimated by biphasic partitioning to *γ*^*LW*^, *γ*^−^, and *γ*^+^ components of the surface tension of particles and phases of ABS. Finally, the surface tension components were used to quantitatively estimate contributions of LRIs in biologically relevant events such as salt-induced nanoparticle agglomeration and protein-nanoparticle interactions using extended DLVO (xDLVO) theory-based calculations.

## Biphasic partitioning of particles

The first objective of this study was to estimate the partitioning coefficient of nanoparticles or biomacromolecules in a biphasic system (**Figure 1-Step 1**). Partitioning of particles in a biphasic system is governed by the interfacial energy^28, 29^. The high interfacial tension of biphasic systems results in the accumulation of particles at the interface with lateral movement of accumulated particles in cases where the interface is curved^30, 31^. Lowering the interfacial tension of biphasic systems results in equilibrium partitioning of particles as a function of both interfacial tension and the particle size^32, 33^. Therefore, we used Aqueous Biphasic Systems (ABS) for the partitioning of nanoparticles and biomacromolecules as ABS have miniscule interfacial tension and are suitable for partitioning of particles up to 100 nm^33^. We used ABS prepared using neutral polymers for partitioning as they do not have an interfacial potential difference and partitioning of particles is not due to their surface charge in such ABS^28^. For partitioning studies, quasi-spherical citrate-capped gold nanoparticles (AuNP) with anionic surface characteristics were synthesized using a modified citrate-reduction method (Figure S1). We also studied the partitioning of Bovine Serum Albumin (BSA) and Lysozyme which are colloidal biomacromolecules. To study the efficacy of the developed framework in estimating the changes in the surface of particles, we cationized BSA to change its surface characteristics without altering the secondary structure content (Figure S2). As all phases of the ABS had different pathlengths for the same suspended volume in a multi-well plate, therefore, a pathlength correction factor was used for accurate measurement of absorbance (Figure S3). For calculating the partitioning coefficient of particles in ABS, we assumed that all nanoparticles and proteins in aqueous suspension had similar surface characteristics and particles partitioned between top and bottom phases (Figure S3). The partitioning coefficient is related to ∆*γ* which is the difference in the interfacial tension between particles and both phases of ABS (Supplementary Section 1)^28^. As the partitioning coefficient is a function of both surface area and surface characteristics of the nanoparticles^28^, the ∆*γ* value obtained for a nanoparticle or a protein in a single ABS is an indicator of its surface characteristics.

## Relationship between ∆γ and surface tension components

The next step was to use ∆*γ* values to estimate particle surface tension (**Figure 1-Step 3**). The partitioning coefficients of cells in ABS with different interfacial tension are related to their contact angle^34^ and therefore can be used to estimate their surface tension components. We developed a theoretical framework for correlating surface tension components of particles to their ∆*γ* value which is calculated using experimentally measured biphasic partitioning data (Supplementary Section 1). The estimated surface tension components were dispersive 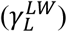, polar 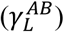, electron donor 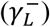, and electron acceptor 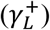 components and should not be considered equivalent to the free energies of the surface. The developed theoretical framework estimates surface tension of particles in as-prepared aqueous state and therefore is not amiable to artifacts introduced by the drying during sample preparation. Changing the dispersion state of colloids, especially proteins, is known to influence the surface tension estimated using contact angle measurement^35, 36^.

For successful implementation of the developed theoretical framework, we needed to resolve the surface tension of top and bottom phases of ABS into *γ*^*LW*^, *γ*^*AB*^, *γ*^−^, and *γ*^+^ components (**Figure 1-Step 2**). We proposed an experimental methodology based on surface tension estimation, contact angle measurement on thin polyethylene (PE) films, and solvatochromic analysis for resolving the surface tension of polymeric phases. We validated the proposed method using water as a model solvent. We measured *γ*_*L*_ of water by the pendant drop method and resolved it into 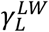 and 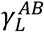 components by measuring the contact angle of water on thin PE films as PE is an apolar polymer^37^. We used the heat-press method to prepare PE films to avoid changes in the physicochemical properties caused by residual solvents left after solvent-casting^38^. Further, there are no reported methods available in literature to resolve 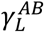 into 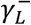 and 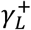 components of surface tension. The 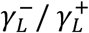 ratio for water proposed in OCG theory is 1:1 while this ratio should be 1:2.343 as per Abraham’s scale of acidity and basicity ^27, 39, 40^. We hypothesized that the Kamlet-Taft’s solvent parameters could be used to determine the 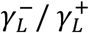 ratio^41^ as these parameters have a linear relationship with partitioning coefficient of proteins^42^. Further, the 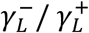 ratio obtained using Kamlet-Taft parameters should be squared in order to get a ratio at par with Abraham’s scale^40, 43^. The 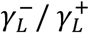 ratio of water estimated by solvatochromic analysis was 2.69 which is in close agreement with Abraham’s scale (Figure S4 and Table S1). The increased acidity of water observed in our measurements is primarily due to the dissolution of atmospheric carbon dioxide^44^. We used the proposed methodology to experimentally estimate *γ*^*LW*^, *γ* ^*AB*^, *γ*^−^, and *γ*^+^ components of the surface tension of polymeric phases of ABS (Table S2).

## Surface tension estimation of colloids by biphasic partitioning

We used the developed theoretical framework (Supplementary Section 1) to estimate *γ*^*LW*^, *γ*^*AB*^, *γ*^−^, and *γ*^+^ components of the surface tension of AuNP and proteins (**Figure 1-Step 3**). We performed a non-linear regression using the biphasic partitioning data from three different ABS (Figure S3).

**Table 1** shows all the values of estimated surface tension components of AuNP and proteins. The surface tension of lysozyme estimated by biphasic partitioning approached the reported values for hydrated lysozyme estimated using contact angle measurement thereby validating the developed method. The resolved components of the surface tension of lysozyme were however different from the reported values as we experimentally estimated the 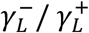 ratios of the phases of ABS. We also evaluated the sensitivity of the developed method in estimating changes in the protein surface. For this, Bovine Serum Albumin (BSA) was cationized (cBSA) by increasing the surface amine group density (Figure S2). Surface basicity estimated as electron donor component (*γ*^−^) correlated well with the changes in the surface amine group density of the proteins estimated by o-phthaldialdehyde (OPA) assay (**Figure 2A**). We observed a linear relationship between the *γ*^−^ of proteins and relative concentration of surface amine groups (**Figure 2B**). The Pearson’s correlation coefficient for the correlation was 0.9999, and the slope was significantly different from zero with a p-value of 0.007 estimated by a 2-tailed t-test.

**Figure 2.**
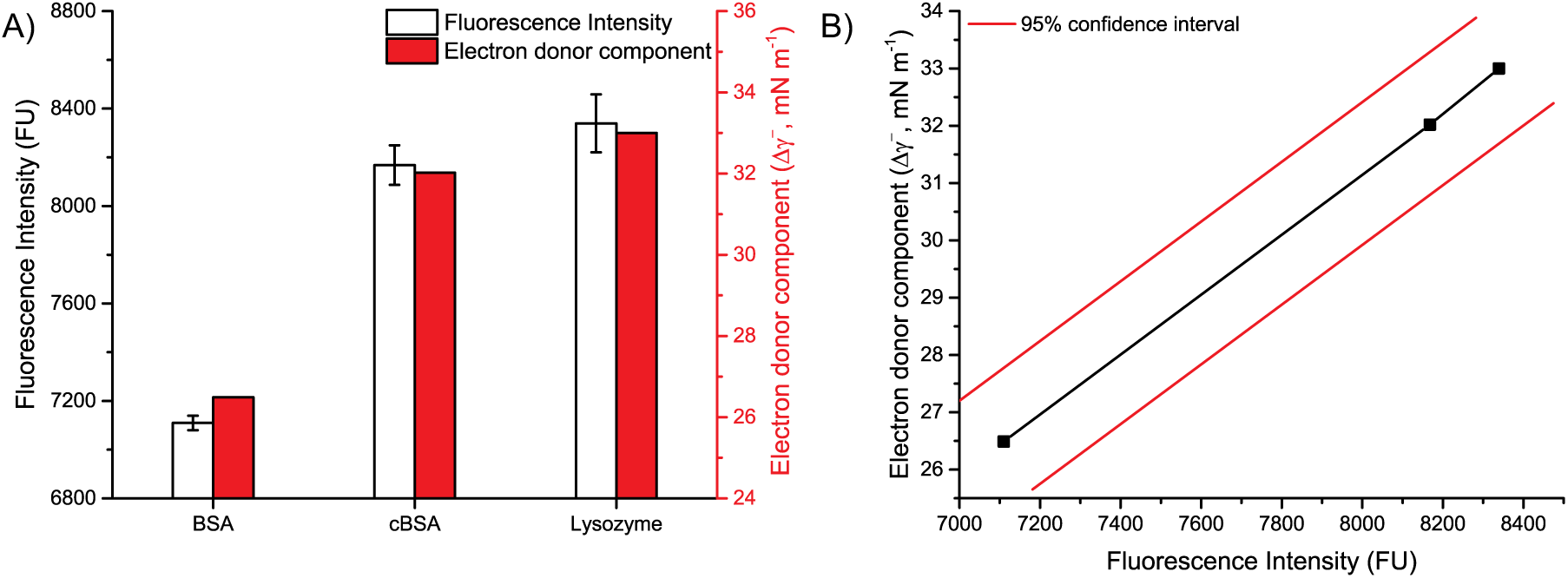
Relationship between surface tension of proteins and their surface chemistry. A) The relative concentration of amine groups on the surface of protein molecule estimated by o-phthaldialdehyde (OPA) assay is indicated by the fluorescence intensity on the left-hand axis. The electron donor component (*γ*^−^) of protein estimated are shown on the right-hand axis. *γ*^−^ is determined by the concentration of amine groups present on the protein surface. B) Correlation between *γ*^−^ and OPA fluorescence.

**Table 1.** Surface tension components estimated by aqueous-biphasic partitioning of aqueous suspensions of gold nanoparticles and model proteins in a fully hydrated state.

|  | Surface tension (mN m<sup>-1</sup>) |  |  |  |  |
| --- | --- | --- | --- | --- | --- |
| | $\gamma$ | $\gamma^{LW}$ | $\gamma^{AB}$ | $\gamma^+$ | $\gamma^-$ |
| Gold nanoparticles (AuNP) | 60.64 | 45.73 | 14.91 | 0.58 | 25.70 |
| Bovine Serum Albumin (BSA) | 58.24 | 45.57 | 12.67 | 1.52 | 26.49 |
| Cationic BSA (cBSA) | 70.58 | 53.61 | 16.97 | 2.25 | 32.02 |
| Lysozyme | 71.79 | 55.59 | 16.21 | 1.99 | 33.00 |
| Lysozyme* (dry)<sup>45</sup> | 48.8 | 41.2 | 2.6 | 0.07 | 23.4 |
| Lysozyme* (hydrated)<sup>45</sup> | ≈72.2 | 31.5 | ≈40.7 | ≈4.5 | ≈56.2 |
| $\gamma$ - surface tension; $\gamma^{LW}$ - Lifshitz-van der Waals component; $\gamma^{AB}$ - acid-base component;<br>$\gamma^+$ - electron acceptor component; $\gamma^-$ - electron donor component; and * represents values reported in the literature | | | | | |

We are for the first time reporting an *in-situ* estimation of the surface tension of hydrated nanoparticles and proteins. The only other reported method used the empirical estimation of the contribution of different LRIs in establishing the nanoparticle-protein interface^16, 17^. The developed method, however, is limited by the size of nanoparticles that can be efficiently partitioned as larger sized particles undergo sedimentation and cannot be partitionied succesfully. Moreover, the surface roughness of nanoparticles also influences the partitioning of nanoparticles in a biphasic system due to contact-line pinning of nanoparticles at the liquid-liquid interface^46, 47^. Therefore, the developed method needs further validation for the cases mentioned above.

The developed method is an ensemble measurement method and therefore estimates the average value of surface tension in a given population of particles having similar surface characteristics. The surface tension components reported here, therefore, do not quantify the chemical heterogeneity on the surface of a single nanoparticle or biomacromolecule. To draw a parallel, zeta potential measurement by electrophoretic mobility is also an ensemble measurement and does not provide any information about the heterogeneity of the surface charge density on the surface of an individual nanoparticle or a protein molecule^48^. The zeta potential, however, can explain how a population of nanoparticles or biomacromolecule will behave at a macroscale, for example, their stability under a given set of conditions. Thus, we hypothesized that average surface characteristics of a nanoparticle or biomacromolecule can help in predicting their population-wide behavior. To test this hypothesis, we further studied the role of LRIs in biologically relevant processes.

## Quantitative estimation of LRIs at the nanoscale

The interplay of two competing phenomena which are the salt-induced agglomeration of nanoparticles and adsorption of biomolecules determine the final state of nanomaterials in the biological milieu and nanoparticle-mediated biological effect^49, 50, 51, 52^. Therefore, we further studied the role of LRIs in governing these biologically relevant processes to gain a mechanistic insight into the nano-bio interfaces. However, the main hindrance for the present work was non-availability of theories which could quantify LRIs at the nanoscale. xDLVO is one of the theories which provides experimentally validated trends for both hydrophobic attraction and hydrophilic repulsion between colloids and a polymeric membrane^53^. Although the xDLVO theory provides information about interfacial EL, LW, and AB interactions for macroscopic homogenous systems^37^, it has been successfully used in estimating the fundamental nature of the interaction between planar surfaces and highly heterogeneous biological samples like proteins^13^ or bacteria^54^. The xDLVO theory-based calculations, however, cannot explain the stochasticity in case a pair of bacteria with significantly different surfaces is used^55^. Although the xDLVO theory is limited in the information it can provide, it can be used for quantitative estimation of LRIs between colloids having similar surface heterogeneity. We, therefore, used xDLVO calculations only for quantitative estimation of LRIs between nanoparticles or nanoparticles and a single protein species.

## Role of LRIs in salt-induced AuNP agglomeration

The agglomeration of nanoparticles upon exposure to biological milieu is a consequence of an interplay between attractive and repulsive LRIs between nanoparticles. We used xDLVO theory (Supplementary Section 2) to calculate the interaction energy between AuNP suspended in water (**Figure 1-Step 3**). Further, we also studied the pH-dependence of salt-induced AuNP agglomeration by calculating interaction energies at varying pH and inter-particle distances equivalent to Debye length corresponding to the ionic strength of the medium. We observed that there was minimal change in the energetics of salt-induced AuNP agglomeration at pH 3-11 (**Figure 3A**). The interaction between AuNP was unfavorable at all pH values at lower ionic strength, whereas, it was favorable at higher ionic strength. We also observed that EL interaction and contribution from Brownian motion (BM) were unfavorable at higher ionic strength, whereas, contributions from LW and AB interactions were favorable (ΔG-ratio plots in **Figure 3A**). The high ionic strength of the medium effectively screens the EL repulsion, and this allows LW and AB interactions to govern the formation of AuNP-AuNP interface. The interaction energy at pH 7.0 as a function of the ionic strength of the medium became increasingly favorable with an increase in NaCl concentration (**Figure 3B**). The contribution of different interaction energies showed a transition at around 25 mM NaCl concentration (**Figure 3A**). These results predicted 25-30 mM as the concentration of NaCl beyond which AuNP should be unstable. For experimental validation, we studied the stability of AuNP in NaCl solution, and we observed AuNP were highly unstable at ionic concentrations higher than 30 mM (**Figure 3**C). A distinct biphasic behavior was observed on either side of the 25-30 mM NaCl concentration with a stable AuNP suspension at low ionic strength conditions even at extended time points and a rapid agglomeration of AuNP at high ionic strength conditions at the onset. We also calculated the non-electrostatic interaction energy (NEIE) between AuNP suspended in water to understand the nature of LRIs at AuNP-AuNP interface (Supplementary Section 3). The overall NEIE was favorable (Figure S5) with a significant contribution from the apolar interaction. Moreover, the AuNP-AuNP and water-water polar cohesion energies were enough to compensate for the polar adhesion between AuNP and water molecules.

**Figure 3.**
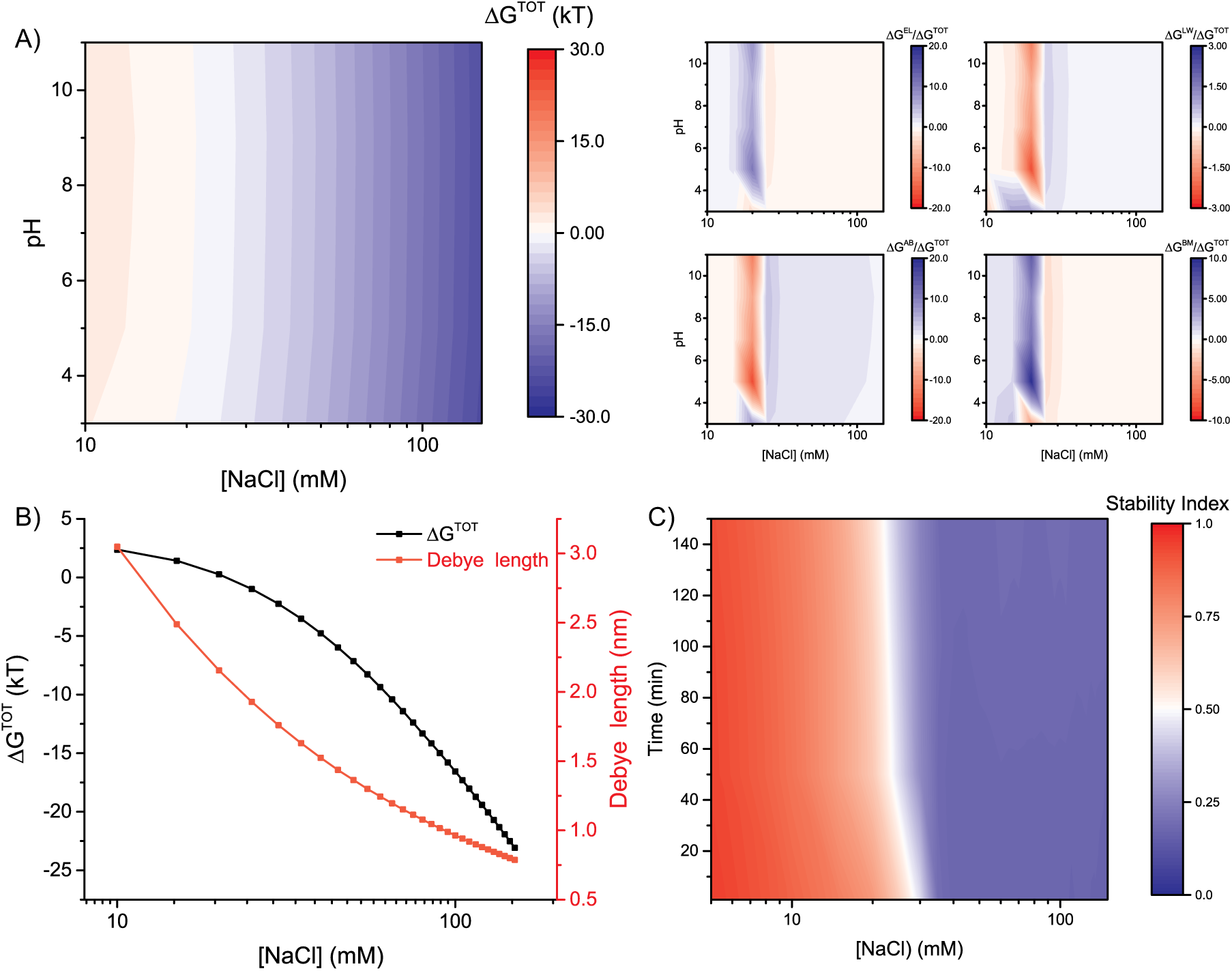
Salt-induced agglomeration of gold nanoparticles (AuNP). A) Interaction energy map of AuNP-AuNP interaction. The interaction energies were calculated at different pH values using xDLVO theory. The interparticle distances for calculations was equal to the Debye length corresponding to the ionic strength of the media. Alongside images show ratio plots of different interaction energies to total interaction energy. The independent contributions from Electrostatic (EL), Lifshitz-van der Waals (LW), and acid-base (AB) interaction energies along with contribution from Brownian Motion (BM) of AuNP are shown. The interpretation of the plot is as follows: red region on interaction energy plot (IE^RED^) and corresponding red region in ratio plot (Ratio^RED^) indicates that LRIs are energetically favorable, however, the interaction is energetically unfavorable. IE^BLUE^ and Ratio^RED^ indicates that LRIs are energetically unfavorable and the interaction is favorable. IE^RED^ and Ratio^BLUE^ indicates both the contribution of LRIs and interaction are energetically unfavorable. IE^BLUE^ and Ratio^BLUE^ indicates both the contribution of LRIs and interaction are energetically favorable. B) Interaction energy calculated at an inter-particle distance equal to Debye length corresponding to the ionic strength of the medium at pH 7.0; C) Stability of AuNP in increasing concentration of NaCl at 1, 50, 150 and 200 min. The stability index values range from 0 (unstable) to 1 (stable).

Collectively, the results presented herein demonstrate that AuNP are relatively stable at low ionic strength environments as EL repulsion provides an energy barrier. As the salt concentration increases, the magnitude of EL repulsion keeps decreasing and the contribution from AB component increases (Figure S5). Therefore, the higher ionic strength of media induces a favorable interaction between AuNP. The AuNP interface is governed majorly by the apolar cohesion between AuNP. Due to this, AuNP once agglomerated cannot be re-dispersed as the contribution from the energy of polar adhesion between AuNP and water is insufficient to overcome the apolar cohesion between AuNP. The data presented here resolves the contribution and interplay between EL, LW, AB, and BM components of interaction energy in salt-induced agglomeration of AuNP. The results corroborate with the experimental observations validating estimated surface tension components using biphasic partitioning.

## Role of LRIs in nanoparticle-protein interactions

Protein adsorption at a liquid-solid interface is a multi-phase process which includes a) approach of protein to a surface, b) adsorption of protein, c) structural rearrangement or denaturation of protein, d) desorption of protein, e) renaturation of protein in some cases^56^. Along the same lines, we envisioned two level of interactions for protein adsorption on a nanoparticle surface as defined for protein-microparticle interactions^13^. The primary interactions due to the approach of proteins to a nanoparticle surface are predominantly interfacial interactions between protein and nanoparticle surface and governed by the surface characteristics of both proteins and nanoparticles. The secondary interactions involve structural rearrangements of the protein due to their adsorption onto the nanoparticle surface and are governed primarily by the structural stability of proteins and the strength of primary interaction. We hypothesized that in the case of nanoparticle-protein interactions, LRIs being most dominant of primary interactions could energetically explain the extent of structural rearrangement of proteins caused by secondary interactions. To test this hypothesis, we used a set of model proteins (BSA and cBSA) which had a similar secondary structure but different surface characteristics (Figure S2). We will not discuss desorption and renaturation of proteins from nanoparticle surface here as it is beyond the scope of this manuscript. However, as a general rule, stable proteins show a higher rate of desorption as they do not undergo structural rearrangements upon adsorption thereby having minimal secondary interactions^57^.

Computational methods have indicated that although the surface of protein molecules is heterogeneous, they prefer a particular orientation during interaction with the nanoparticle surface due to residual-specific interactions^58^. Therefore, we used xDLVO theory to estimate the average interfacial interaction energy between model proteins and nanoparticles as a function of pH of the media and inter-particle distance equivalent to the Debye length (Supplementary Section 2). The estimated interfacial energies may not accurately represent LRIs, but they provide information about their importance in forming the nanoparticle-protein interface. The pH and ionic strength dependent interaction energy maps of AuNP with proteins revealed interaction energies are unfavorable at low ionic strengths, whereas, they are favorable at higher ionic strengths (**Figure 4**). In case of AuNP-BSA interaction (**Figure 4**A), we observed that at low ionic strength conditions, the change in contribution from various interactions was maximum around the isoelectric point of BSA (pH = 5.0). Contributions from BM were unfavorable at all pH values whereas, contributions from EL were unfavorable above the isoelectric point of BSA. In case of LW and AB interactions, the contribution is favorable at higher ionic strengths and unfavorable at low ionic strength. In case of AuNP-cBSA interaction (**Figure 4**B), the contribution from EL, LW, and AB interaction was favorable, whereas, the contribution from BM was unfavorable at higher ionic strength at all pH values. The contribution from the interaction energy was highest for AB interaction for both BSA and cBSA indicating a significant influence of aqueous environment in driving the AuNP-protein interaction. In literature, there are instances where protein-nanoparticle interactions are studied at low ionic strength conditions as nanoparticles are unstable at ionic strengths corresponding to physiological conditions.^59^ However, our results indicate that using low ionic strength media in estimating nanoparticle-protein interactions can lead to erroneous interpretation about biological behavior of nanoparticles. As there are no experimental methods available for quantification of primary or secondary protein-nanoparticle interactions, therefore, we could not validate the results predicted by xDLVO theory.

**Figure 4.**
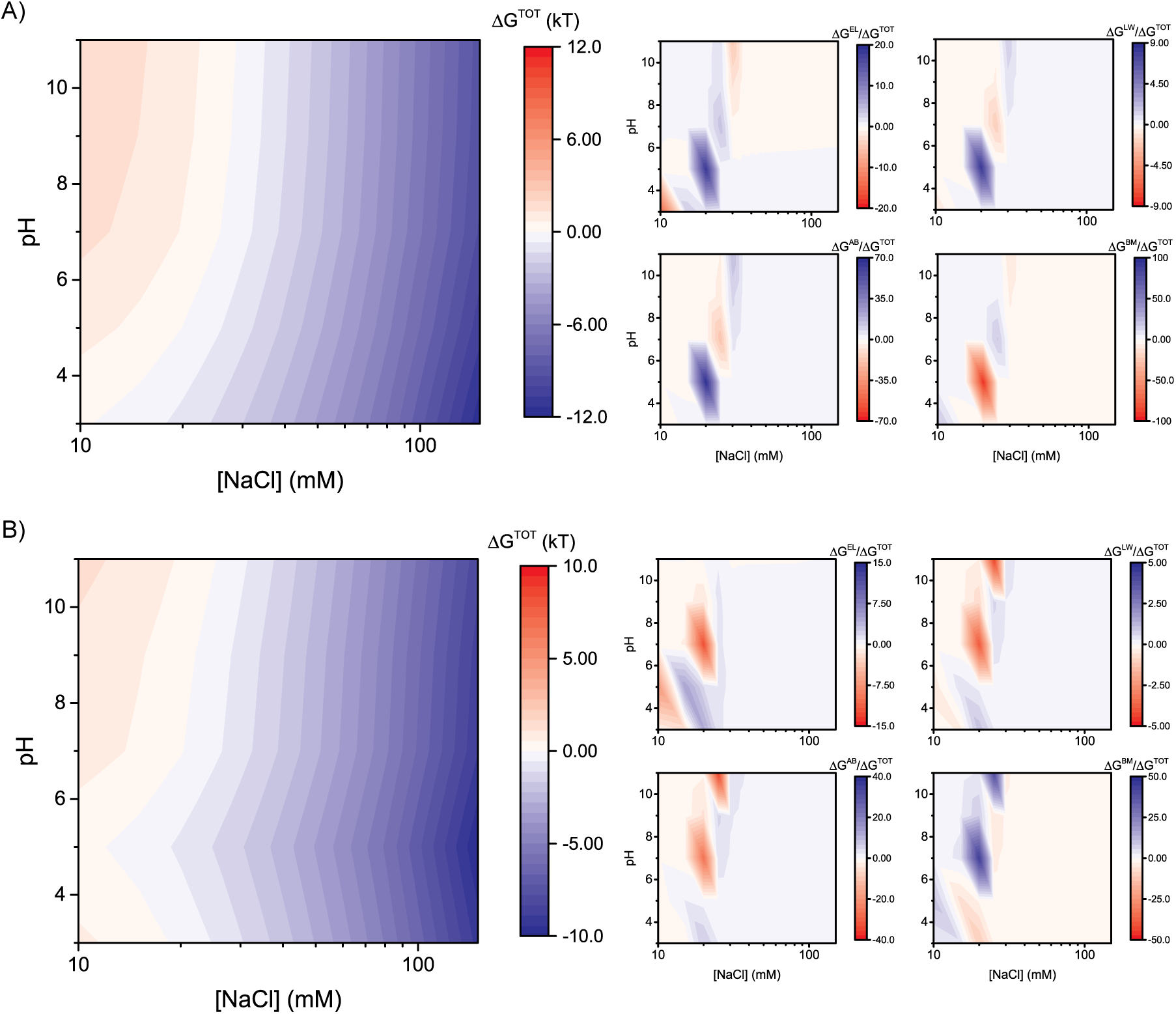
Primary interaction between gold nanoparticles (AuNP) and proteins. Interaction energy maps for A) AuNP-BSA interaction and B) AuNP-cBSA interaction. The interaction energies were calculated at different pH values using xDLVO theory. The interparticle distances for calculations was equal to the Debye length corresponding to the ionic strength of the media. Alongside images show ratio plots of different interaction energies to total interaction energy. The independent contributions from Electrostatic (EL), Lifshitz-van der Waals (LW), and acid-base (AB) interaction energies along with contribution from Brownian Motion (BM) of AuNP are shown. The interpretation of the plot is as follows: red region on interaction energy plot (IE^RED^) and corresponding red region in ratio plot (Ratio^RED^) indicates that LRIs are energetically favorable, however, the interaction is energetically unfavorable. IE^BLUE^ and Ratio^RED^ indicates that LRIs are energetically unfavorable and the interaction is favorable. IE^RED^ and Ratio^BLUE^ indicates both the contribution of LRIs and interaction are energetically unfavorable. IE^BLUE^ and Ratio^BLUE^ indicates both the contribution of LRIs and interaction are energetically favorable.

We also calculated NEIE between both AuNP-BSA and AuNP-cBSA (Supplementary Section 1 and Figure S6). The overall NEIE is favorable and comparable for interaction of AuNP with both BSA and cBSA, indicating that EL interactions have a deterministic role in establishing the AuNP-protein interface in this case. The apolar adhesion between proteins and AuNP was more favorable as compared to apolar adhesion between water and proteins/nanoparticles indicating that once the favorable interaction takes place, desorption of BSA and cBSA will be minimal. Although not observed for the AuNP, it is reported in literature that BSA molecules once undergone structural rearrangements after adsorption on the anionic silica nanoparticles do not desorb from the surface^60^.

Further, we also studied the structural integrity of protein in presence and absence of nanoparticles using circular dichroism (CD) spectroscopy. The CD spectra of BSA and cBSA in presence and absence of AuNP was collected after 24 hours of incubation and the curves were fitted to a reference dataset on DichroWeb using a CONTIN algorithm^61^. Figure S7 shows two dimensional plots of change in ellipticity of proteins as a function of wavelength and temperature in presence and absence of AuNP. We observed that there was no significant difference in the secondary structure of native proteins after 24 hours of incubation (**Figure 5** and Table S3). However, in presence of AuNP, we observed a statistically significant difference in the content of helices, sheets, and unordered structures of cBSA, whereas, in case of BSA we only observed a significant change in unordered structures (**Figure 5** and Table S3). Moreover, temperature alone significantly explained the variance of the data in heat-melting of BSA in presence of AuNP, whereas, for cBSA the effect of both temperature and presence of AuNP influenced the variance of the data (Table S4). Therefore, we concluded that the interaction between cBSA and AuNP perturbed the secondary structure of cBSA. Our results are consistent with reports of BSA adsorption on planar surfaces which have demonstrated an increase in the β-sheet content at the expense of helical content.^62^ Moreover, the increase in the β-sheet content in case of adsorption of peptides on planar surfaces is primarily due to an interplay between the hydrophobic and electrostatic interactions.^63^

We further studied the thermodynamics of structural rearrangements of proteins in presence of AuNP by studying protein melting using changes in the ellipticity at 222 nm as a function of temperature.^64^ We observed a significant change in the enthalpy, entropy, and melting temperature of cBSA in the presence of AuNP, whereas, no such changes were observed for BSA (Figure S7). We also calculated the difference in the energy required for the denaturation of native proteins in presence or absence of AuNP. We observed that in case of cBSA there was a significant energy difference indicating a loss of secondary structure as a result of interaction with nanoparticles. This indicated that due to the loss in secondary structure, cBSA required a lesser amount of heat for denaturation in the presence of AuNP. To correlate xDLVO calculations and CD spectroscopy data, we estimated the free energy of melting required for BSA and cBSA in presence and absence of AuNP at 25 °C for 10 mM NaCl concentration (**Table 2**). The primary interaction energy calculated between AuNP and proteins also indicated that under the studied conditions, the interaction between cBSA-AuNP was favorable, whereas, that between AuNP-BSA was unfavorable. The primary interaction energy was able to compensate for the energy required for the unfolding of cBSA. Therefore, cBSA showed a change in the secondary structure on interacting with AuNP, whereas, the secondary structure of BSA remained unaffected indicated that EL interaction played a significant role in primary interaction energies. Taken together, these results demonstrate the significant role played by LRIs at the nano-bio interface in both binding and structural alteration of adsorbed proteins.

**Figure 5.**
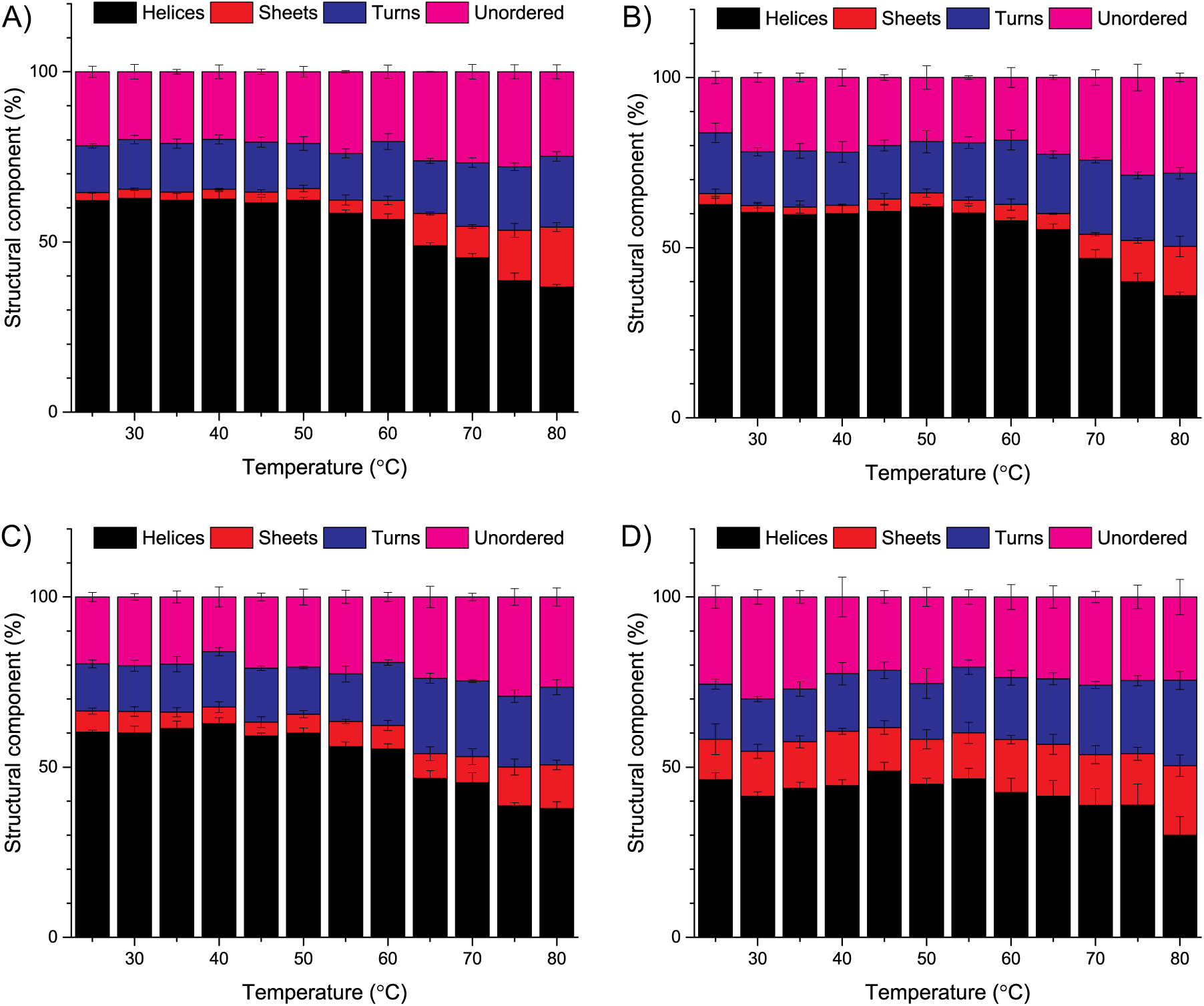
Temperature dependent changes in the secondary structure elements of proteins in native and gold nanoparticle (AuNP) bound state. The percentage of secondary structure elements at different temperatures was estimated by fitting the circular dichroism data using DichroWeb. The percentage secondary structure components in terms of helices, sheets, turns and unordered structure are reported for A) BSA; B) BSA in presence of AuNP; C) cBSA; and D) cBSA in presence of AuNP. The error bars represent sample standard deviation for three independent measurements.

**Table 2.** Free energy required for the denaturation of BSA and cBSA at 25 °C.

| Sample | $\Delta G_m$ | $\Delta\Delta G_m$ | $\Delta G_{int}$ |
| --- | --- | --- | --- |
| BSA | 2.80 | - | - |
| BSA + AuNP | 2.73 | 0.063 | 0.022<br>(EL = 0.794; LW = -0.163; AB = -0.609) |
| cBSA | 2.13 | - | - |
| cBSA + AuNP | 0.86 | 1.279 | -1.675<br>(EL = -1.003; LW = -0.164; AB = -0.508) |
| $\Delta G_m$ is the Gibbs free energy required for protein unfolding at 25 °C in kJ mol<sup>-1</sup> calculated using CD spectroscopy data.; $\Delta\Delta G_m$ – Difference in Gibbs free energy required for protein unfolding between native and AuNP associated proteins in kJ mol<sup>-1</sup>, $\Delta G_{int}$ – Primary interaction energy between protein and AuNP calculated using xDLVO theory for pH 7.0 and ionic strength of 10 mM NaCl. (EL- electrostatic interaction, LW-Lifshitz-van der Waals interaction, AB – acid-base interaction). | | | |

## Conclusions

We have developed a framework based on biphasic partitioning which enabled us to estimate surface tension components of nanoparticles and biomacromolecules suspended in an aqueous suspension. We also developed an experimental framework for estimating the interfacial environment of each phase of ABS and correlated it to the biphasic partitioning. The developed framework is suitable for deciphering the role of LRIs in establishing a nano-bio interface. The experimental validations strengthen the potential of the developed framework to predict the biological behavior of nanoparticles. Moreover, the developed framework is simple, inexpensive, and does not require high-end equipment or extensive sample preparation.

However, there remains a need to develop new biphasic systems for partitioning of larger nanoparticles to increase the scope of the work. The lack of theories for interfacial interactions at the nanoscale is the biggest hindrance to developing an interfacial interaction-based fundamental understanding of nano-bio interfaces. Although, the xDLVO theory provided experimentally verifiable results for the interplay between LRIs at nanoscale interactions, however, it only enables the study of only simple interactions either between nanoparticles or nanoparticle and single protein. The xDLVO theory cannot explain the stochastic information about the interaction of nanoparticles with multiple proteins which is essential for developing a fundamental understanding about the biomolecular corona formation. Collectively, results presented in this manuscript highlight the importance of the developed framework in predicting the biological behavior of nanoparticles. Therefore, the developed framework has a far-reaching impact on nanomaterial screening for biomedical applications as it might facilitate the selection of nanomaterials for biomedical applications by identifying nanomaterials causing adverse biological effects due to their propensity to adsorb certain biomacromolecules.

## Supporting information

## Associated Content

### Supporting Information

The supporting information consist of experimental methods and other procedural details, supplementary data in form of figures and tables, and supplementary sections on (1) mathematical model of estimation of nanoparticle surface tension using biphasic partitioning in ABS, (2) mathematical model for xDLVO calculations, and (3) equations for non-electrostatic interaction energy calculations.

### Author Contributions

VS and DSK conceived the project. VS designed the experiments, performed the research and analyzed the results. SD performed the nanoparticle-protein interaction studies and VS and SD analyzed the results for the same. All authors discussed the results and contributed equally to the writing of the paper.

## Acknowledgements

The work presented in this manuscript was carried out using the research grant given by Nano Mission, Department of Science and Technology, Govt. of India under grant no. SR/NM/NS-1006/2010. VS and SD acknowledge the financial assistance provided by MHRD, Govt. of India. Authors acknowledge Prof. Krishnacharya for providing access to goniometer for surface tension and contact angle measurements and Advanced Imaging, IIT Kanpur for Transmission Electron Microscope facility. Authors also acknowledge the infrastructural support provided by IIT Kanpur during the research.

## Reference

1. Nel AE, Mädler L, Velegol D, Xia T, Hoek EMV, Somasundaran P, et al. Understanding Biophysicochemical Interactions at The Nano-Bio Interface. Nat Mater 2009, 8(7): 543–557.

2. Tenzer S, Docter D, Kuharev J, Musyanovych A, Fetz V, Hecht R, et al. Rapid Formation of Plasma Protein Corona Critically Affects Nanoparticle Pathophysiology. Nat Nanotechnol 2013, 8(10): 772–781.

3. Chithrani BD, Ghazani AA, Chan WCW. Determining the Size and Shape Dependence of Gold Nanoparticle Uptake into Mammalian Cells. Nano Lett 2006, 6(4): 662–668.

4. Walkey CD, Olsen JB, Song F, Liu R, Guo H, Olsen DWH, et al. Protein Corona Fingerprinting Predicts the Cellular Interaction of Gold and Silver Nanoparticles. ACS Nano 2014, 8(3): 2439–2455.

5. Jiang W, Kim BYS, Rutka JT, Chan WCW. Nanoparticle-Mediated Cellular Response is Size-Dependent. Nat Nanotechnol 2008, 3(3): 145–150.

6. Deng ZJ, Liang M, Monteiro M, Toth I, Minchin RF. Nanoparticle-Induced Unfolding of Fibrinogen Promotes Mac-1 Receptor Activation and Inflammation. Nat Nanotechnol 2011, 6(1): 39–44.

7. Schöttler S, Becker G, Winzen S, Steinbach T, Mohr K, Landfester K, et al. Protein Adsorption is Required for Stealth Effect of Poly(ethylene glycol)-and Poly(phosphoester)-Coated Nanocarriers. Nat Nanotechnol 2016, 11(4): 372–377.

8. Bos R, Van Der Mei HC, Busscher HJ. Physico-Chemistry of Initial Microbial Adhesive Interactions - Its Mechanisms and Methods for Study. FEMS Microbiol Rev 1999, 23(2): 179–229.

9. van Oss CJ, Docoslis A, Wu W, Giese RF. Influence of macroscopic and microscopic interactions on kinetic rate constants. I. Role of the extended DLVO theory in determining the kinetic adsorption constant of proteins in aqueous media, using von Smoluchowski’s approach. Colloids Surf, B 1999, 14(1-4): 99–104.

10. Škvarla J. A Physico-chemical Model of Microbial Adhesion. J Chem Soc, Faraday Trans 1993, 89(15): 2913–2921.

11. Van Oss CJ. The Forces Involved in Bioadhesion to Flat Surfaces and Particles – Their Determination and Relative Roles. Biofouling 1991, 4(1-3): 25–35.

12. French RH, Parsegian VA, Podgornik R, Rajter RF, Jagota A, Luo J, et al. Long Range Interactions in Nanoscale Science. Rev Mod Phys 2010, 82(2): 1887–1944.

13. van Oss CJ, Docoslis A, Giese Jr RF. Free Energies of Protein Adsorption onto Mineral Particles - From the Initial Encounter to the Onset of Hysteresis. Colloids Surf, B 2001, 22(4): 285–300.

14. Gessner A, Lieske A, Paulke BR, Müller RH. Influence of surface charge density on protein adsorption on polymeric nanoparticles: Analysis by two-dimensional electrophoresis. Eur J Pharm Biopharm 2002, 54(2): 165–170.

15. Shang L, Yang L, Seiter J, Heinle M, Brenner-Weiss G, Gerthsen D, et al. Nanoparticles Interacting with Proteins and Cells: A Systematic Study of Protein Surface Charge Effects. Advanced Materials Interfaces 2014, 1(2): 1300079.

16. Xia XR, Monteiro-Riviere NA, Riviere JE. An Index for Characterization of Nanomaterials in Biological Systems. Nat Nanotechnol 2010, 5(9): 671–675.

17. Xia XR, Monteiro-Riviere NA, Mathur S, Song X, Xiao L, Oldenberg SJ, et al. Mapping the Surface Adsorption Forces of Nanomaterials in Biological Systems. ACS Nano 2011, 5(11): 9074–9081.

18. van Oss CJ, Ju L, Chaudhury MK, Good RJ. Estimation of the Polar Parameters of the Surface Tension of Liquids by Contact Angle Measurements on Gels. J Colloid Interface Sci 1989, 128(2): 313–319.

19. JonΑš A, Karadag Y, Tasaltin N, Kucukkara I, Kiraz A. Probing Microscopic Wetting Properties of Superhydrophobic Surfaces by Vibrated Micrometer-Sized Droplets. Langmuir 2011, 27(6): 2150–2154.

20. Yu J, Wang H, Liu X. Direct Measurement of Macro Contact Angles Through Atomic Force Microscopy. Int J Heat Mass Transfer 2013, 57(1): 299–303.

21. Sundberg M, Månsson A, Tågerud S. Contact Angle Measurements by Confocal Microscopy for Non-Destructive Microscale Surface Characterization. J Colloid Interface Sci 2007, 313(2): 454–460.

22. Brugnara M, Volpe CD, Siboni S, Zeni D. Contact Angle Analysis on Polymethylmethacrylate and Commercial Wax by Using an Environmental Scanning Electron Microscope. Scanning 2006, 28(5): 267–273.

23. Weon BM, Lee JS, Kim JT, Pyo J, Je JH. Colloidal Wettability Probed with X-ray Microscopy. Curr Opin Colloid Interface Sci 2012, 17(6): 388–395.

24. Isa L, Lucas F, Wepf R, Reimhult E. Measuring Single-Nanoparticle Wetting Properties by Freeze-Fracture Shadow-Casting Cryo-Scanning Electron Microscopy. Nat Commun 2011, 2: 438.

25. Leo A, Hansch C, Elkins D. Partition Coefficients and Their Uses. Chem Rev 1971, 71(6): 525–616.

26. Abbott NJ. Prediction of Blood–brain Barrier Permeation in Drug Discovery From *In Vivo*, *In Vitro* And *In Silico Models*. Drug Discovery Today: Technol 2004, 1(4): 407–416.

27. Praetorius A, Tufenkji N, Goss K-U, Scheringer M, von der Kammer F, Elimelech M. The Road to Nowhere: Equilibrium Partition Coefficients for Nanoparticles. Environ Sci: Nano 2014, 1(4): 317–323.

28. Albertsson PA. Partition of Cell Particles and Macromolecules: Distribution and Fractionation of Cells, Viruses, Microsomes, Proteins, Nucleic Acids, and Antigen-Antibody Complexes in Aqueous Polymer Two-Phase Systems. J. Wiley, 1960.

29. Dasgupta S, Auth T, Gompper G. Nano-and Microparticles at Fluid and Biological Interfaces. J Phys: Condens Matter 2017, 29(37): 373003.

30. Hristovski KD, Westerhoff PK, Posner JD. Octanol-Water Distribution of Engineered Nanomaterials. J Environ Sci Health, Part A: Toxic/Hazard Subst Environ Eng 2011, 46(6): 636–647.

31. Sharifi-Mood N, Liu IB, Stebe KJ. Curvature Capillary Migration of Microspheres. Soft Matter 2015, 11(34): 6768–6779.

32. Helfrich MR, El-Kouedi M, Etherton MR, Keating CD. Partitioning and Assembly of Metal Particles and Their Bioconjugates in Aqueous Two-Phase Systems. Langmuir 2005, 21(18): 8478–8486.

33. Luechau F, Ling TC, Lyddiatt A. A Descriptive Model and Methods for Up-Scaled Process Routes for Interfacial Partition of Bioparticles in Aqueous Two-Phase Systems. Biochem Eng J 2010, 50(3): 122–130.

34. Gerson DF. Cell surface energy, contact angles and phase partition. I. Lymphocytic cell lines in biphasic aqueous mixtures. Biochim Biophys Acta 1980, 602(2): 269–280.

35. van Oss CJ, Absolom DR, Neumann AW, Zingg W. Determination of The Surface Tension of Proteins I. Surface Tension of Native Serum Proteins in Aqueous Media. Biochim Biophys Acta, Protein Struct 1981, 670(1): 64–73.

36. Absolom DR, van Oss CJ, Zingg W, Neumann AW. Determination of Surface Tensions of Proteins II. Surface Tension of Serum Albumin, Altered at the Protein-Air Interface. Biochim Biophys Acta, Protein Struct 1981, 670(1): 74–78.

37. van Oss CJ, Chaudhury MK, Good RJ. Interfacial Lifshitz-van der Waals and Polar Interactions in Macroscopic Systems. Chem Rev 1988, 88(6): 927–941.

38. Briscoe B, Smith A. Rheology of Solvent-Cast Polymer Films. J Appl Polym Sci 1983, 28(12): 3827–3848.

39. Abraham MH. Scales of Solute Hydrogen-Bonding: Their Construction and Application to Physicochemical and Biochemical Processes. Chem Soc Rev 1993, 22(2): 73–83.

40. Della Volpe C, Siboni S. Some Reflections on Acid–Base Solid Surface Free Energy Theories. J Colloid Interface Sci 1997, 195(1): 121–136.

41. Marcus Y. The Properties of Organic Liquids That are Relevant to Their Use as Solvating Solvents. Chem Soc Rev 1993, 22(6): 409–416.

42. Madeira PP, Reis CA, Rodrigues AE, Mikheeva LM, Chait A, Zaslavsky BY. Solvent Properties Governing Protein Partitioning in Polymer/Polymer Aqueous Two-Phase Systems. J Chromatogr A 2011, 1218(10): 1379–1384.

43. Della Volpe C, Siboni S. Acid–Base Surface Free Energies of Solids and the Definition of Scales in the Good–van Oss–Chaudhury Theory. J Adhes Sci Technol 2000, 14(2): 235–272.

44. Duan Z, Sun R. An Improved Model Calculating CO_2_ Solubility in Pure Water and Aqueous NaCl Solutions From 273 to 533 K and From 0 to 2000 Bar. Chem Geol 2003, 193(3-4): 257–271.

45. van Oss CJ. The Properties of Water and their Role in Colloidal and Biological Systems. Academic Press, 2008.

46. Manoharan VN. Colloids at Interfaces: Pinned Down. Nat Mater 2015, 14(9): 869.

47. Wang A, McGorty R, Kaz DM, Manoharan VN. Contact-Line Pinning Controls How Quickly Colloidal Particles Equilibrate with Liquid Interfaces. Soft Matter 2016, 12(43): 8958–8967.

48. Sikora A, Bartczak D, Geißler D, Kestens V, Roebben G, Ramaye Y, et al. A systematic comparison of different techniques to determine the zeta potential of silica nanoparticles in biological medium. Analytical Methods 2015, 7(23): 9835–9843.

49. Cedervall T, Lynch I, Lindman S, Berggård T, Thulin E, Nilsson H, et al. Understanding the Nanoparticle-Protein Corona Using Methods to Quantify Exchange Rates and Affinities of Proteins for Nanoparticles. Proc Natl Acad Sci U S A 2007, 104(7): 2050–2055.

50. Cedervall T, Lynch I, Foy M, Berggård T, Donnelly SC, Cagney G, et al. Detailed Identification of Plasma Proteins Adsorbed on Copolymer Nanoparticles. Angew Chem Int Ed 2007, 46(30): 5754–5756.

51. Walczyk D, Bombelli FB, Monopoli MP, Lynch I, Dawson KA. What the cell “sees” in bionanoscience. J Am Chem Soc 2010, 132(16): 5761–5768.

52. Huang D, Zhou H, Liu H, Gao J. The Cytotoxicity of Gold Nanoparticles is Dispersity-Dependent. Dalton Trans 2015, 44(41): 17911–17915.

53. Brant JA, Childress AE. Membrane-Colloid Interactions: Comparison of Extended DLVO Predictions with AFM Force Measurements. Environ Eng Sci 2002, 19(6): 413–427.

54. Bayoudh S, Othmane A, Mora L, Ben Ouada H. Assessing Bacterial Adhesion Using DLVO and XDLVO Theories and the Jet Impingement Technique. Colloids Surf, B 2009, 73(1): 1–9.

55. Chia TWR, Nguyen VT, McMeekin T, Fegan N, Dykes GA. Stochasticity of Bacterial Attachment and Its Predictability by the Extended Derjaguin-Landau-Verwey-Overbeek Theory. Appl Environ Microbiol 2011, 77(11): 3757–3764.

56. Norde W. Adsorption of (Bio)Polymers, with Special Emphasis on Globular Proteins. Colloids and Interfaces in Life Sciences and Bionanotechnology. CRC Press, 2011, pp 277–304.

57. Karlsson M, Ekeroth J, Elwing H, Carlsson U. Reduction of Irreversible Protein Adsorption on Solid Surfaces by Protein Engineering for Increased Stability. J Biol Chem 2005, 280(27): 25558–25564.

58. Ding F, Radic S, Chen R, Chen P, Geitner NK, Brown JM, et al. Direct Observation of a Single Nanoparticle–Ubiquitin Corona Formation. Nanoscale 2013, 5(19): 9162–9169.

59. Huang R, Carney RP, Stellacci F, Lau BL. Protein–Nanoparticle Interactions: The Effects of Surface Compositional and Structural Heterogeneity are Scale Dependent. Nanoscale 2013, 5(15): 6928–6935.

60. Satzer P, Svec F, Sekot G, Jungbauer A. Protein Adsorption onto Nanoparticles Induces Conformational Changes: Particle Size Dependency, Kinetics, and Mechanisms. Eng Life Sci 2016, 16(3): 238–246.

61. Whitmore L, Wallace BA. Protein Secondary Structure Analyses from Circular Dichroism Spectroscopy: Methods and Reference Databases. Biopolymers 2008, 89(5): 392–400.

62. Norde W, Giacomelli CE. BSA Structural Changes During Homomolecular Exchange Between the Adsorbed and the Dissolved States. J Biotechnol 2000, 79(3): 259–268.

63. Giacomelli CE, Norde W. Conformational Changes of the Amyloid β-Peptide (1–40) Adsorbed on Solid Surfaces. Macromol Biosci 2005, 5(5): 401–407.

64. Greenfield NJ. Using Circular Dichroism Collected as a Function of Temperature to Determine the Thermodynamics of Protein Unfolding and Binding Interactions. Nat Protoc 2006, 1(6): 2527–2535.

