## Supplementary material for "Quantitative Estimation of Long-Range Interactions at the Nanoscale"

**Supporting information includes following sections**


| | |
|---|---|
| Supplementary Figures | S3 |
| Supplementary Tables | S9 |
| Supplementary Section 1 | S13 |
| Supplementary Section 2 | S16 |
| Supplementary Section 3 | S19 |
| Supplementary Methods | S21 |
| Supplementary References | S31 |

### Supplementary Figures

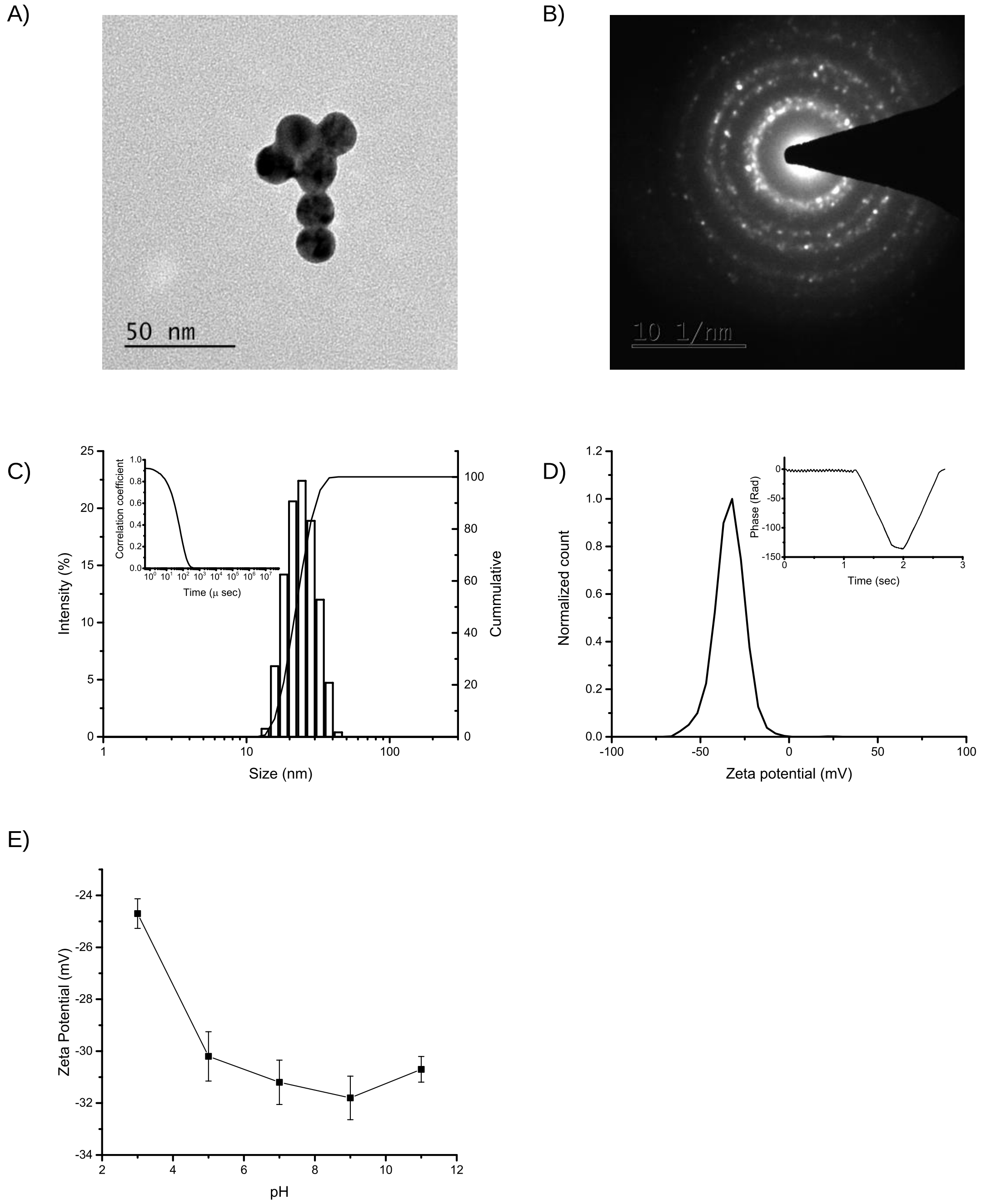


**Figure S1**. Characterization of gold nanoparticles (AuNP) synthesized using citrate reduction method. A) Transmission electron micrograph of AuNP showing their quasi-spherical morphology (Scale bar - 50 nm); B) Small area diffraction pattern of AuNP showing polycrystalline nature of AuNP; C) Frequency distribution of Z-average (left-hand axis) and cumulative frequency (right-hand axis) of AuNP. The inset shows intensity decay correlogram showing the absence of aggregates in the AuNP suspension; D) Zeta potential distribution of AuNP. The inset shows phase plot for the same sample; and E) Change in the zeta potential of AuNP as a result of a change in the pH of the surrounding medium.

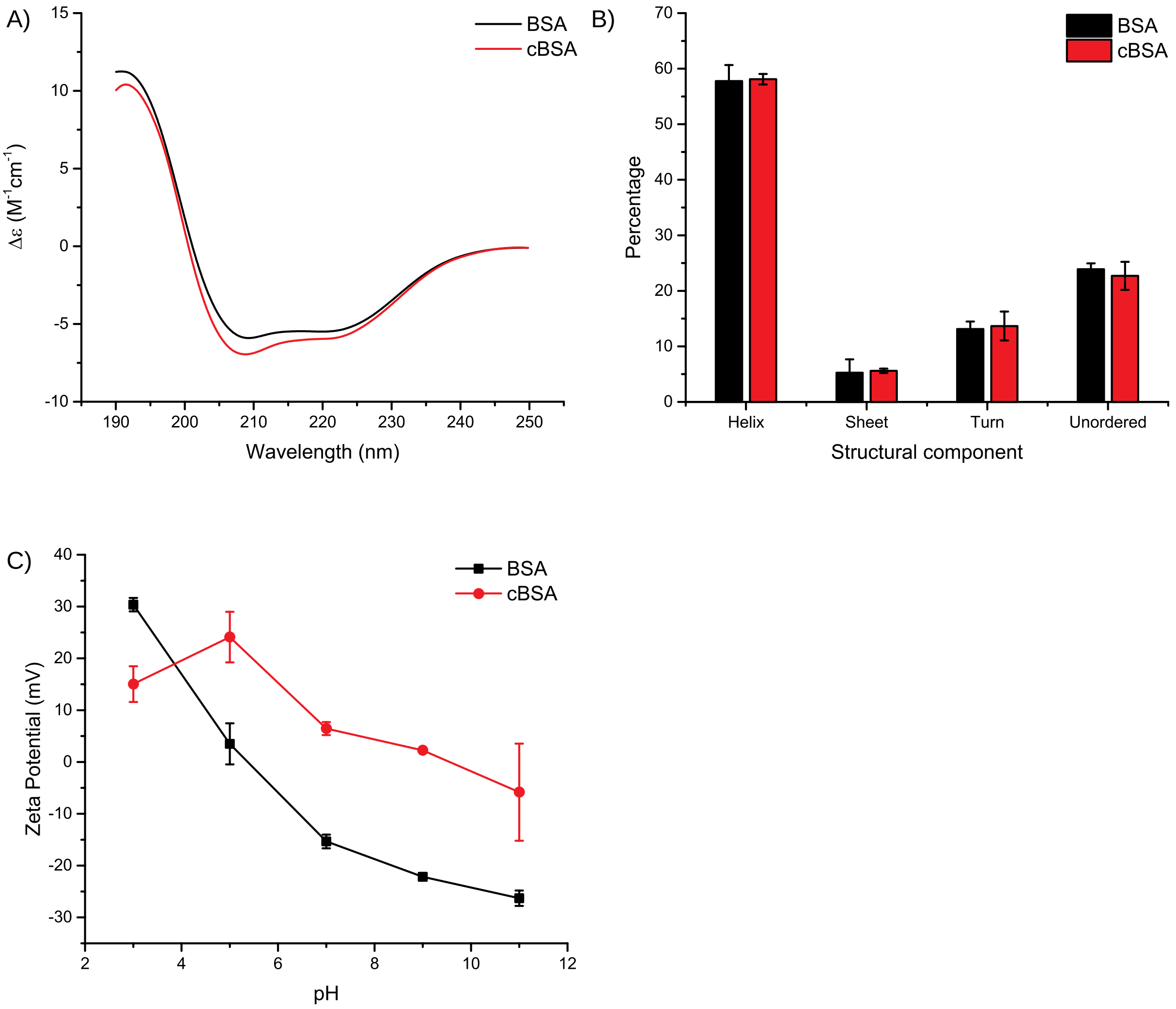


**Figure S2**. Protein modification and characterization. A) CD spectra, and B) percentage of secondary structure components of BSA and cationic BSA (cBSA). C) Change in the zeta potential of BSA and cBSA as a result of a change in the pH of the surrounding medium.

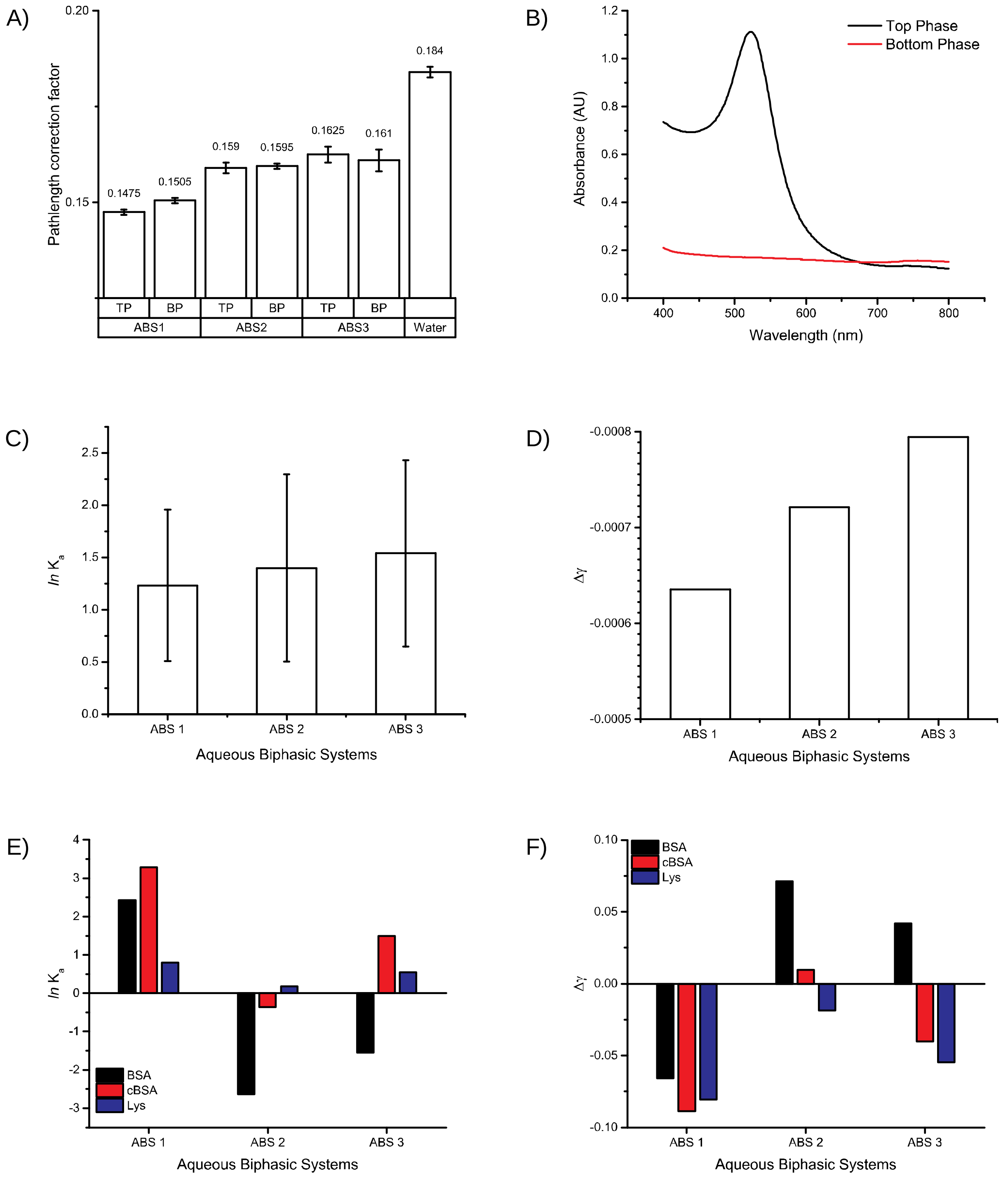


**Figure S3**. Partitioning of gold nanoparticles (AuNP) and model proteins in Aqueous Biphasic Systems (ABS). ABS1 was a biphasic system of PEG 1000 and Dextran 40000; ABS 2 was a biphasic system of PEG 6000 and Dextran 40000; and ABS3 was a biphasic system of PEG 10000 and Dextran 40000. A) Pathlength correction factors for all the phases of ABS; B) UV-vis absorbance spectra of AuNP in top and bottom phases of ABS showing preferential partitioning of AuNP in the top phase; C) Partition coefficient of AuNP in ABS; and D) Δγ values of AuNP calculated using the mean values of partition coefficient; E) Natural log of the partition coefficient of model proteins in ABS. BSA – bovine serum albumin, cBSA – cationized BSA, and LYS- lysozyme. Average values of six samples were taken for the calculation. F) Δγ values of model proteins calculated using the mean values of partition coefficient.

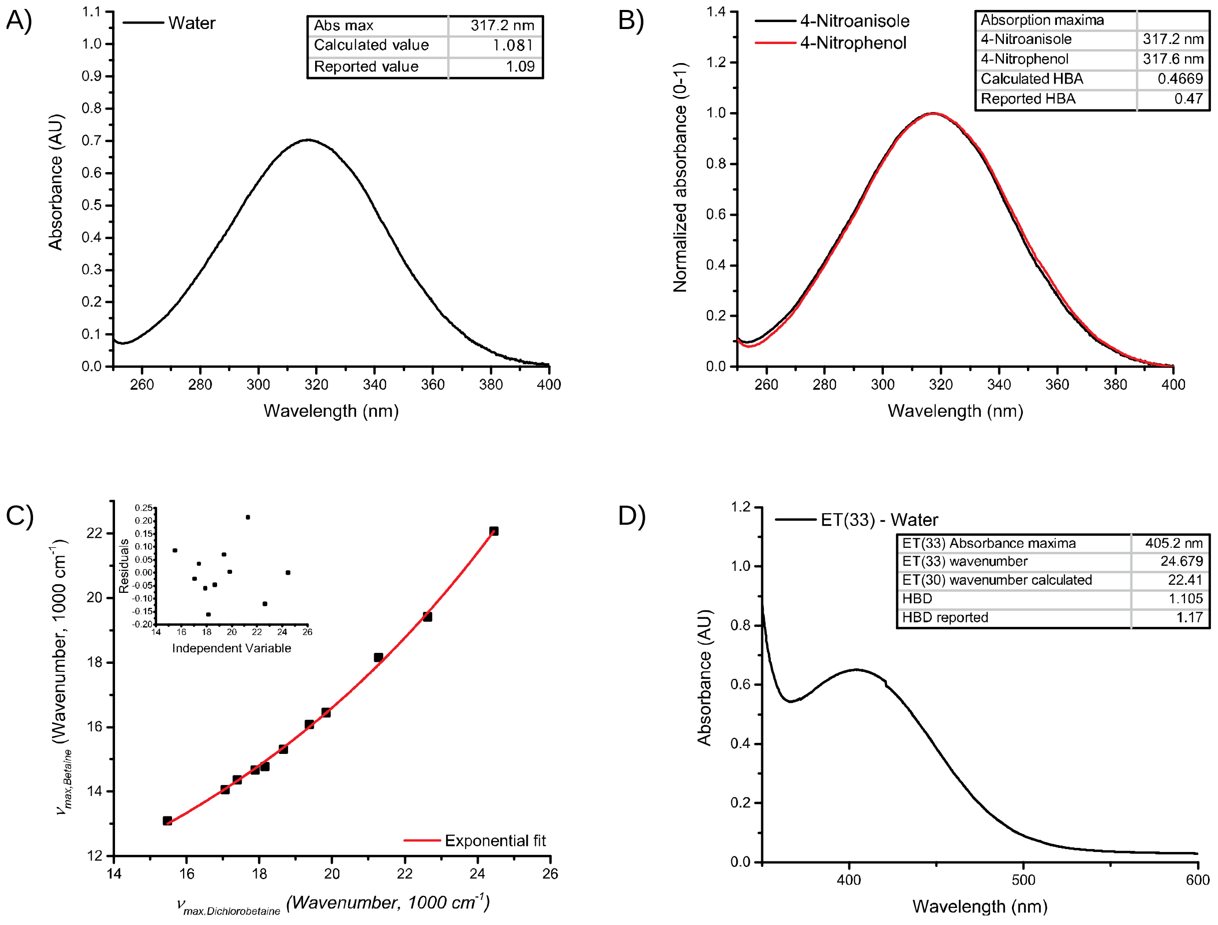


**Figure S4**. Resolution of the surface tension of water into different components using solvatochromic analysis. A) Polarizability (π*); B) Hydrogen bond acceptance/electron pair donation ability to form a coordinative bond (β); C) Relationship between wavenumber of absorbance maxima of betaine and dichlorobetaine for different solvents. An exponential fit was performed, and the inset shows residual values for the fit, and D) Hydrogen bond donation ability (α).

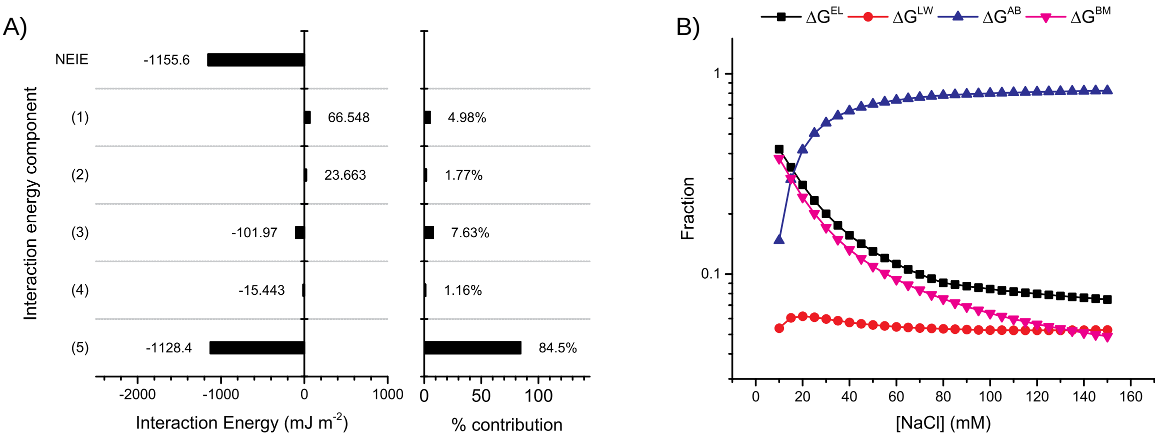


**Figure S5.** A) Contribution and percentage contribution of different interaction energy components towards non-electrostatic interaction energy (NEIE) between AuNP suspended in water. The interaction energy components are as follows: (1) polar adhesive energy between electron-acceptors of water molecules and electron-donors of the AuNP surface; (2) polar adhesive energy between electron-acceptors of AuNP surface and electron-donors of water molecules; (3) polar cohesive energy between electron-acceptors and electron-donors of water molecules; (4) polar cohesive energy between electron-acceptors and electron-donors of the AuNP surface; (5) apolar adhesive (Lifshitz-van der Waals) interaction energy between AuNP and water molecules. B) The fraction of different interaction energies between gold nanoparticles (AuNP) as a function of the ionic strength of the media at pH 7.0

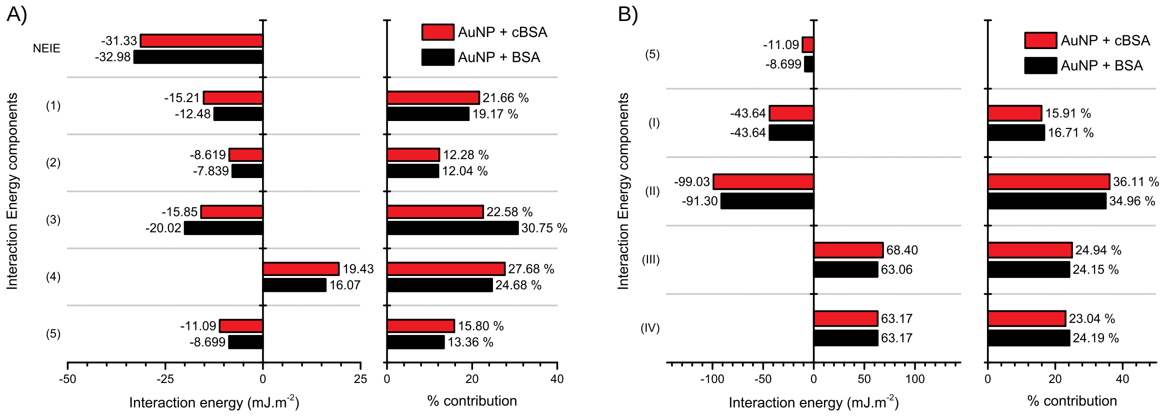


**Figure S6.** A) Contribution and percentage contribution of different components of interaction energy towards non-electrostatic interaction energy (NEIE) between AuNP and proteins. Different components are as follows: (1) Polar adhesion between electron donor groups of AuNP surface tension and electron acceptor groups of protein surface tension; (2) polar adhesion between electron acceptor groups of AuNP surface tension and electron donor groups of protein surface tension; (3) polar adhesion between electron acceptor groups of AuNP and proteins and electron donor ability of water molecules; (4) polar adhesion between electron donor groups of AuNP and proteins and electron acceptor ability of water molecules; (5) apolar interaction between AuNP and protein. B) The apolar component further constitutes different interactions as follows: (I) apolar cohesion between water molecules; (II) apolar adhesion between AuNP and proteins; (III) apolar adhesion between protein and water molecules; and (IV) apolar adhesion between AuNP and water molecules.

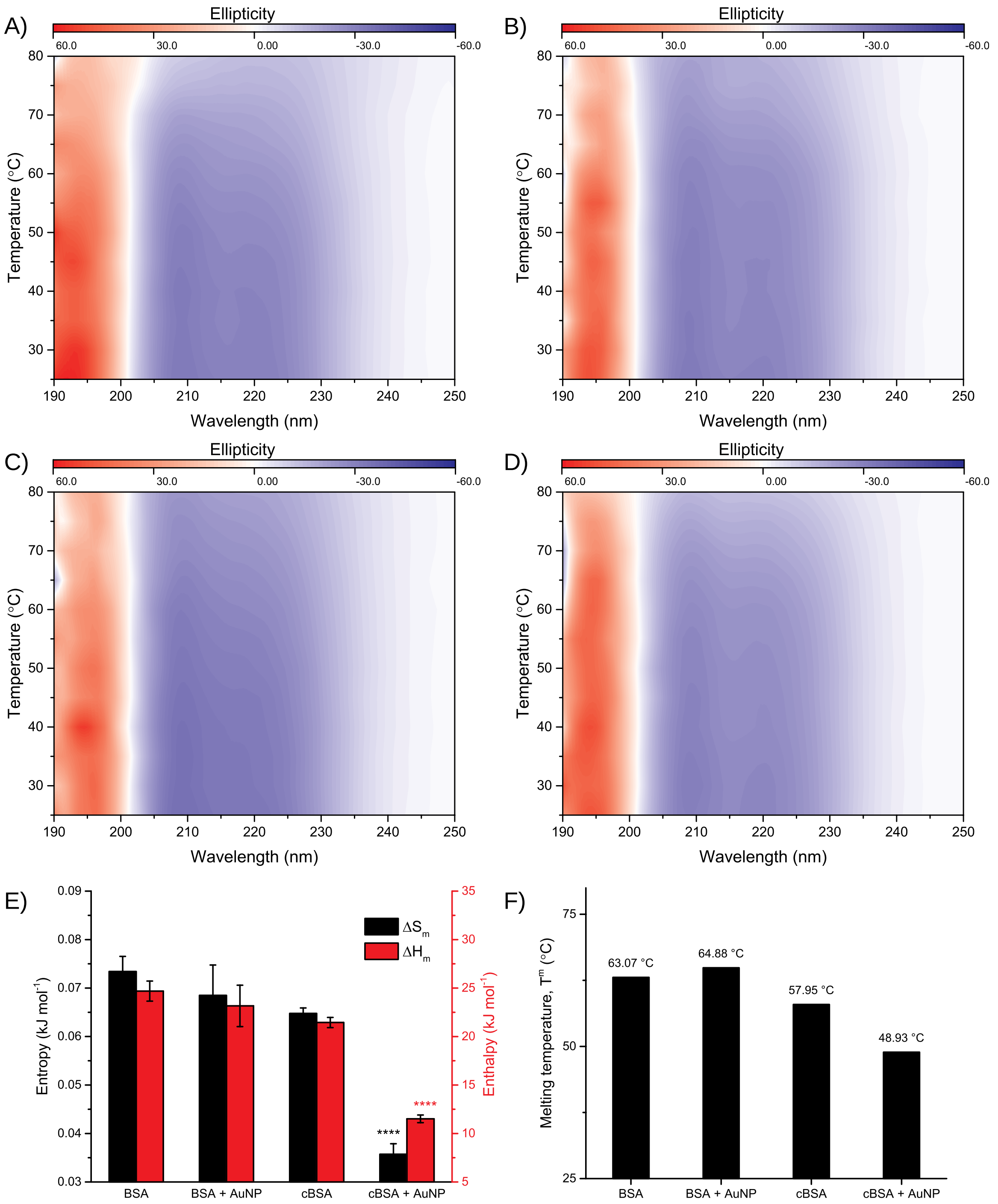


**Figure S7**. Structural changes in adsorbed protein after their interaction with gold nanoparticles (AuNP). Two-dimensional CD spectra of A) BSA; B) BSA in presence of AuNP; C) cBSA; and D) cBSA in presence of AuNP. For protein-AuNP interaction, AuNP were incubated with respective proteins in as-synthesized state followed by 24 hours incubation at 25 °C. E) Thermodynamic parameters of protein melting in presence and absence of AuNP. Enthalpic and entropic contributions for the temperature dependent protein denaturation estimated using the temperature-dependent changes in the ellipticity values of proteins at 222 nm. The significance was tested using Ordinary one-way ANOVA with Holm-Sidak's multiple comparison test. The comparison was done between BSA vs BSA+AuNP and cBSA vs cBSA+AuNP. **** means p-value less than 0.001. F) Melting temperature of proteins in the native state and in presence of AuNP.

### Supplementary Tables

**Table S1.** Resolved surface tension components of water

| **Quantity** | **Measured** | **Reported[1]** |
|---|---|---|
| Surface tension of water ($\gamma$), mJ.m$^{-2}$ | - | 72.8 |
| Contact angle of water on LDPE | 102.8 | - |
| Surface tension of LDPE, mJ.m$^{-2}$ | - | 36.8 |
| $\gamma^{LW}$ component of surface tension of LDPE, mJ.m$^{-2}$ | - | 36.8 |
| $\gamma^{LW}$component of surface tension of water ($\gamma^{LW}$), mJ.m$^{-2}$ | 21.818 | 21.8 |
| $\gamma^{AB}$ component of surface tension of water ($\gamma^{AB}$), mJ.m$^{-2}$ | 50.982 | 51.0 |
| Acidity to basicity ratio | 2.367 | 2.343[2] |
| Basic component of surface tension of water ($\gamma^{-}$), mJ.m$^{-2}$ | 10.77 | 25.5 |
| Acidic component of surface tension of water ($\gamma^{+}$), mJ.m$^{-2}$ | 60.34 | 25.5 |

**Table S2.** Resolved surface tension components of top and bottom phases of ABS

| ABS | Phase | $\alpha$ | $\beta$ | $\pi^*$ | $x$ | $\gamma_L$ | $\theta_{PE}$ | $\gamma_L^{LW}$ | $\gamma_L^{AB}$ | $\gamma_L^-$ | $\gamma_L^+$ |
|---|---|---|---|---|---|---|---|---|---|---|---|
| Water | - | 1.08 | 0.48 | 1.29 | 2.69 | 72.8 | 102.8 | 21.82 | 50.98 | 10.77 | 60.34 |
| PEG 1: Dex 40 | Top phase | 1.1 | 0.51 | 1.16 | 2.27 | 64.6 | 81.83 | 38.12 | 26.48 | 5.83 | 30.07 |
| | Bottom phase | 1.12 | 0.49 | 1.15 | 2.35 | 63.98 | 84.37 | 34.57 | 29.41 | 6.26 | 34.55 |
| PEG 6: Dex 40 | Top phase | 1.1 | 0.49 | 1.16 | 2.37 | 63.02 | 84.97 | 32.91 | 30.11 | 6.36 | 35.67 |
| | Bottom phase | 1.14 | 0.55 | 1.24 | 2.28 | 62.72 | 84.33 | 33.26 | 29.46 | 6.47 | 33.52 |
| PEG 10: Dex 40 | Top phase | 1.15 | 0.41 | 1.17 | 2.89 | 63.12 | 86.97 | 30.93 | 32.19 | 5.56 | 46.59 |
| | Bottom phase | 1.17 | 0.43 | 1.13 | 2.64 | 63.59 | 84.8 | 33.69 | 29.91 | 5.66 | 39.53 |

**Footnote:** $\alpha$ – Hydrogen bond donation (HBD); $\beta$ – Hydrogen bond acceptance (HBA) or electron pair donation ability to form a coordinative bond; $\pi^*$ – Polarity/ polarizability parameter; $x$ – ratio between HBD and HBA parameters ($\alpha/\beta$); $\gamma_L$ – surface tension of phase; $\theta_{PE}$ – contact angle of phase on LDPE thin film (in degrees); $\gamma_L^{LW}$ – Lifshitz – van der Waals component of surface energy of phase; $\gamma_L^{AB}$ – Acid-base component of surface energy of phase; $\gamma_L^-$ – electron donor component of phase; $\gamma_L^+$ – electron acceptor component of phase. (all $\gamma$ values are in $mN.m^{-1}$). ABS1 was a biphasic system of PEG 1000 and Dextran 40000; ABS 2 was a biphasic system of PEG 6000 and Dextran 40000; and ABS3 was a biphasic system of PEG 10000 and Dextran 40000.

**Table S3.** Ordinary two-way ANOVA analysis with Holm-Sidak multiple comparison test for studying the changes in secondary structure of proteins after 24 hours of incubation. The CD spectra were recorded at 25 °C (n=3). P-values are reported for each comparison values in bold represent statistically significant difference.

| **Comparisons** | | | | **Helices** | **Sheets** | **Turns** | **Unordered** | **Significance** |
|---|---|---|---|---|---|---|---|---|
| **BSA** | **BSA + AuNP** | **cBSA** | **cBSA + AuNP** | | | | | |
| * | * | | | 0.7854 | 0.5867 | 0.1236 | **0.0112** | Effect of AuNP on BSA |
| * | | * | | 0.4699 | 0.0866 | 0.9380 | 0.2256 | Comparison between proteins |
| * | | | * | **<0.0001** | **<0.0001** | 0.5066 | **0.0866** | |
| | * | * | | 0.4351 | 0.1792 | 0.1236 | 0.1051 | |
| | * | | * | **<0.0001** | **<0.0001** | 0.5566 | **<0.0001** | Comparison between effect of AuNP on proteins |
| | | * | * | **<0.0001** | **0.0082** | 0.5066 | **0.0067** | Effect of AuNP on cBSA |

**Table S4**. Ordinary Two-way ANOVA analysis with Holm-Sidak multiple comparison test for studying the effect of temperature and AuNP on the secondary structure components of BSA and cBSA as estimated by CD spectroscopy (n=3). The effect of temperature and presence or absence of AuNP was studied for a particular secondary structure content. P-values are reported along with the percentage of variation explained by each variable in parenthesis. Values in bold represent statistically significant difference.

| | **Secondary structure** | **Interaction** | **Temperature** | **AuNP** |
|---|---|---|---|---|
| **BSA vs BSA+AuNP** | Helices | **0.0129 (1.77%)** | **<0.0001 (95.18%)** | 0.5587 (0.022%) |
| | Sheets | **0.0028 (3.41%)** | **<0.0001 (90.5%)** | **0.0005 (1.38%)** |
| | Turns | 0.8542 (3.42%) | **<0.0001 (58.74%)** | **<0.0001 (10.99%)** |
| | Unordered | **0.0024 (13.92%)** | **<0.0001 (65.00%)** | **0.0206 (2.25%)** |
| **cBSA vs cBSA+AuNP** | Helices | **<0.0001 (9.06%)** | **<0.0001 (47.45%)** | **<0.0001 (37.07%)** |
| | Sheets | 0.3278 (3.69%) | **<0.0001 (20.13%)** | **<0.0001 (62.51%)** |
| | Turns | 0.0848 (8.01%) | **<0.0001 (70.06%)** | **0.0225 (2.28%)** |
| | Unordered | **0.0006 (28.43%)** | **0.0008 (27.55%)** | **0.0002 (11.37%)** |

**Supplementary Section 1**

**Nanoparticle partitioning in biphasic systems**

Biphasic systems are characterized by the existence of two phases in thermodynamic equilibrium. Nanoparticles show a diameter-dependent partitioning behavior when they are suspended in a biphasic system.[3] This partitioning behavior is a result of two forces[4]: (i) the Brownian motion of nanoparticles leading to a random distribution of nanoparticles in both phases, and (ii) interplay of interfacial tension between the two phases and nanoparticles resulting in an uneven distribution of nanoparticles between phases. The partition behavior of nanoparticle can be explained by the interfacial tensions between different interfaces (between the nanoparticle surface and two phases). These interfaces include

1. Liquid-liquid interface between the top and bottom phases with $\gamma_{TB}$ as the interfacial tension.
2. Nanoparticle–liquid interface between nanoparticles and the top phase with $\gamma_{NT}$ as the interfacial tension.
3. Nanoparticle–liquid interface between nanoparticles and the bottom phase with $\gamma_{NB}$ as the interfacial tension.

Depending on their affinity to a either top or bottom phase, nanoparticles distribute between different phases of ABS. The nanoparticle distribution in the ABS is a function of the difference between particle interfacial tension in the top and bottom phase ($\Delta\gamma$, equation 1) and diameter of nanoparticles ($D_N$) or surface area of nanoparticles ($A$).[4] Equation 2 shows the relationship between particle distribution in bulk phases and $\Delta\gamma$. Similar equations can be written for particle distribution between top phase and interface (equation 3) or between bottom phase and interface (equation 4). In these equations, $K_{TB}$ is the partitioning coefficient of nanoparticles between the top and bottom phases, $G_{IT}$ is the partitioning coefficient of nanoparticles between the interface and the top phase, $G_{IB}$ is the partitioning coefficient of nanoparticles between the interface and the bottom phase, $k$ is the Boltzmann's constant (1.38 x $10^{-14}$ N.nm.$K^{-1}$ or 1.381 x $10^{-23}$ J.$K^{-1}$), and $T$ is the absolute temperature in Kelvin. These equations provide a relationship between the observable equilibrium phenomena (i.e. partitioning coefficient) and the diameter of nanoparticles. Since $\Delta\gamma$ is a function of nanoparticle surface area, it can be used as an indicator of surface characteristics of nanoparticles. For example, if $\Delta\gamma$ is similar for different sized nanoparticles, then their surface characteristics are identical, and partitioning of these particles is a result of the difference in the surface area. The adsorption of nanoparticles at the interface is a function of the diameter of the particles, whereas, for non-spherical particles, nanoparticle adsorption is a function of the surface area of nanoparticles. To conclude, for the nanoparticles partitioning to the top phase, the partition coefficient will tend to increase ($\gamma_{NT} < \gamma_{NB}$) or decrease ($\gamma_{NT} > \gamma_{NB}$) as the surface area of nanoparticle increases.

$$\Delta\gamma = \gamma_{NT} - \gamma_{NB} \qquad 1$$

$$\ln K_{TB} = \ln\left(\frac{\text{Concentration of AuNP in top phase } (C_T)}{\text{Concentration of AuNP in bottom phase } (C_B)}\right) = -\frac{\pi\, D_N^2\, \Delta\gamma}{kT} = -\frac{A\, \Delta\gamma}{kT} \qquad 2$$

$$\ln G_{IT} = \ln\left(\frac{\text{Number of AuNP adsorbed at interface } (N_{Int})}{\text{Number of AuNP/mL in top phase } (N_T)}\right) = \frac{\pi\, D_N^2\, (\Delta\gamma + \gamma_{TB})^2}{4\,\gamma_{TB}\, kT} = \frac{A\, (\Delta\gamma + \gamma_{TB})^2}{4\,\gamma_{TB}\, kT} \qquad 3$$

$$\ln G_{IB} = \ln\left(\frac{\text{Number of AuNP adsorbed at interface } (N_{Int})}{\text{Number of AuNP/mL in bottom phase } (N_B)}\right) = \frac{\pi\, D_N^2\, (\Delta\gamma - \gamma_{TB})^2}{4\,\gamma_{TB}\, kT} = \frac{A\, (\Delta\gamma - \gamma_{TB})^2}{4\,\gamma_{TB}\, kT} \qquad 4$$

**The relationship between $\Delta\gamma$ and surface tension components of nanoparticles.** $\Delta\gamma$ values obtained by partitioning were used to estimate the surface tension of colloids and further resolve it into dispersive and polar components. There was no report of relationship between $\Delta\gamma$ and surface tension of colloids was available in the literature as in the case of planar surfaces wherein the surface tension can be estimated by measuring the contact angle of a known solvent on that surface. Dupre's and Young-Dupre's equations correlate the contact angle and surface tension of a surface and a liquid to work done as shown in equations 5 and 6.

According to Dupre's equation

$$W_{SL} = \gamma_S + \gamma_L - \gamma_{SL} \qquad 5$$

According to Young-Dupre's equation[5-6]

$$W_{SL} = (1 + \cos\theta)\, \gamma_L \qquad 6$$

The contact angle theory was extended by the OCG theory[7] to include Lifshitz-van der Waals ($\gamma^{LW}$) and acid-base ($\gamma^{AB}$) components of surface tension. The OCG theory correlates an observable phenomenon (i.e. contact angle) to the surface tension characteristics of a surface and a liquid as shown in equation 7. To solve the equation, contact angle is measured with three different liquids as there are three unknown terms in it.

$$(1+\cos\theta)\gamma_L = 2\left(\sqrt{\gamma_S^{LW}\gamma_L^{LW}} + \sqrt{\gamma_S^{+}\gamma_L^{-}} + \sqrt{\gamma_S^{-}\gamma_L^{+}}\right) \qquad 7$$

Equations 5, 6, and 7 can be used to define the interface formed between a colloid and a single phase of an ABS (equation 8).

$$\gamma_S + \gamma_L - \gamma_{SL} = 2\left(\sqrt{\gamma_S^{LW}\gamma_L^{LW}} + \sqrt{\gamma_S^{+}\gamma_L^{-}} + \sqrt{\gamma_S^{-}\gamma_L^{+}}\right) \qquad 8$$

Equation 8 explains the interfacial tension between the nanoparticle and either the top or bottom phases of ABS. Subtracting the bottom phase equation from the top phase equation yielded a generalized expression for nanoparticle partitioning in a biphasic system (equation 9). In the resulting equation, the difference in the interfacial tension of colloids with the top and bottom phases ($\Delta\gamma$) was equal to the polar and dispersive components of the colloids and the phases in which they are getting partitioned.

$$\Delta\gamma = (\gamma_T - \gamma_B) - 2\sqrt{\gamma_N^{LW}}\left(\sqrt{\gamma_T^{LW}} - \sqrt{\gamma_B^{LW}}\right) - 2\sqrt{\gamma_N^{+}}\left(\sqrt{\gamma_T^{-}} - \sqrt{\gamma_B^{-}}\right) - 2\sqrt{\gamma_N^{-}}\left(\sqrt{\gamma_T^{+}} - \sqrt{\gamma_B^{+}}\right) \qquad 9$$

Nanoparticle partitioning yielded $\Delta\gamma$ values based on equation 2. There are three unknown variables in equation 9 – surface acidity/electron acceptor component ($\gamma_N^{+}$), surface basicity/electron donor component ($\gamma_N^{-}$), and surface Lifshitz-van der Waals/dispersive component ($\gamma_N^{LW}$). Therefore, nanoparticle partitioning in three different ABS is required to resolve nanoparticle surface tension into different components. This relationship is the first instance in the literature to reconcile theories of particle partitioning in ABS and van OCG's theory of surface energy.

**Supplementary Section 2**

**The xDLVO theory for estimation of LRIs.** Estimation of the decay of the interaction energy as a function of distance provides an insight into the role of LRIs in establishing an interface. The DLVO theory gives an estimation of total interaction energy ($\Delta G^{TOT}$) which includes contributions from electrostatic interaction energy ($\Delta G^{EL}$) and Lifshitz-van der Waals interaction energy ($\Delta G^{LW}$). The DLVO theory, however, fails to explain the phenomenon of hydrophilic repulsions.[8] Therefore, the DLVO theory has been extended to include contributions from the polar interactions which include acid-base interactions.[9] The xDLVO theory provided results which were comparable to those obtained by atomic force microscopy (AFM) for both hydrophobic attraction and hydrophilic repulsion[8]. According to the xDLVO theory, $\Delta G^{TOT}$ includes contributions from acid-base interaction energy ($\Delta G^{AB}$) and energy contribution due to Brownian motion ($\Delta G^{BM}$) in addition to $\Delta G^{EL}$ and $\Delta G^{LW}$. For $\Delta G^{BM}$, a constant value of 1.5 kT was added to the total. The total interaction energy of nanoparticle interaction ($\Delta G^{TOT}$) is given by equation 10.

$$\Delta G^{TOT} = \Delta G^{EL} + \Delta G^{LW} + \Delta G^{AB} + \Delta G^{BM} \qquad 10$$

The distance dependent decay of different components of interaction energies between colloids of radii r1 and r2 separated by a distance $\ell$ was calculated by the set of equations provided in the next section.

**A) Lifshitz-van der Waals interaction energy**

The decay of Lifshitz-van der Waals interaction energy with separation distance $\ell$ is given by equations 11-14. The Lifshitz-van der Waals interaction energy between two colloids at the distance of closest approach $\ell_o$ is calculated by using equation 15 and $\gamma^{LW}$ components of the surface tension of nanoparticles and proteins was obtained from partitioning data.

$$\Delta G^{LW}_{(\ell)} = -\frac{A}{6}\left[\frac{2\,r_1 r_2}{f1_{(r_1,r_2,\ell)}} + \frac{2\,r_1 r_2}{f2_{(r_1,r_2,\ell)}} + \ln\left(\frac{f1_{(r_1,r_2,\ell)}}{f2_{(r_1,r_2,\ell)}}\right)\right] \qquad 11$$

$$f1_{(r_1,r_2,\ell)} = \ell^2 + 2\,r_1\ell + 2r_2\ell \qquad 12$$

$$f2_{(r_1,r_2,\ell)} = \ell^2 + 2\,r_1\ell + 2r_2\ell + 4r_1r_2 \tag{13}$$

$$A = -12\,\pi\,\ell_\circ^2\,\Delta G_{\ell_\circ}^{LW} \tag{14}$$

$$\Delta G_{\ell_\circ}^{LW} = -2\gamma_M^{LW} - 2\sqrt{\gamma_{c1}^{LW}\gamma_{c2}^{LW}} + 2\sqrt{\gamma_{c1}^{LW}\gamma_M^{LW}} + 2\sqrt{\gamma_{c2}^{LW}\gamma_M^{LW}} \tag{15}$$

Where, $\ell$ is the separation distance between two colloids, $\Delta G_{(\ell)}^{LW}$ is the Lifshitz-van der Waals interaction energy between two colloids at a separation distance $\ell$, A is the Hamakar constant, $r_1$ is the radius of colloid 1, $r_2$ is the radius of colloid 2, $\ell_o$ is the distance of closest approach between colloid 1 and colloid 2, and $\Delta G_{\ell_\circ}^{LW}$ is the Lifshitz-van der Waals interaction energy between colloid 1 and colloid 2 at a separation distance of $\ell_o$.

**B) Lewis acid-base interaction energy**

The decay of Lewis acid-base interaction energy between colloids with separation distance $\ell$ is given by equation 16. $\Delta G_{\ell_\circ}^{AB}$ was calculated using equation 17 and electron donor/ acceptor components of the surface tension of nanoparticles and proteins was estimated by partitioning studies.

$$\Delta G_{(\ell)}^{AB} = \pi\,\lambda\,\Delta G_{\ell_\circ}^{AB}\,\frac{(r_1r_2)}{(r_1+r_2)}\,e^{\left[\frac{(\ell_\circ-\ell)}{\lambda}\right]} \tag{16}$$

$$\Delta G_{\ell_\circ}^{AB} = -2\sqrt{\gamma_{c1}^{+}\gamma_{c2}^{-}} - 2\sqrt{\gamma_{c2}^{+}\gamma_{c1}^{-}} + 2\sqrt{\gamma_{c1}^{+}\gamma_{M}^{-}} + 2\sqrt{\gamma_{M}^{+}\gamma_{c1}^{-}} + 2\sqrt{\gamma_{c2}^{+}\gamma_{M}^{-}} + 2\sqrt{\gamma_{M}^{+}\gamma_{c2}^{-}} - 4\sqrt{\gamma_{M}^{+}\gamma_{M}^{-}} \tag{17}$$

Where, $\ell$ is the separation distance between colloid 1 and colloid 2, $\Delta G_{(\ell)}^{AB}$ is the Lewis acid-base interaction energy between colloid 1 and colloid 2 at a distance $\ell$, $\lambda$ is the correlation length for a colloid in the medium and its value is taken as 0.6 nm, $\Delta G_{\ell_\circ}^{AB}$ is the Lewis acid-base interaction energy between colloid 1 and colloid 2 at a separation distance of $\ell_o$, $r_1$ is the radius of colloid 1, $r_2$ is the radius of colloid 2, and $\gamma^+$ is the acidic/electron acceptor and $\gamma^-$ is the basic/electron donor component of surface tension of colloid 1 (c1), colloid 2 (c2) and medium (M).

**C) Electrostatic interaction energy**

For electrostatic interaction energy calculations, the Bos *et al.* equation with the Sader modification was used (equation 18).[10-11]

$$\Delta G_{(\ell)}^{EL} = \pi\varepsilon\,(\zeta_{c1}^2 + \zeta_{c2}^2)\frac{r_1 r_2}{(r_1 + r_2 + \ell)}\left[\frac{2\,\zeta_{c1}\zeta_{c2}}{(\zeta_{c1}^2 + \zeta_{c2}^2)}\ \ln\frac{1 + e^{-\kappa\ell}}{1 - e^{-\kappa\ell}} + \ln 1 - e^{-2\kappa\ell}\right] \qquad 18$$

Where, $\ell$ is the distance between colloid 1 and colloid 2, $\Delta G_{(\ell)}^{EL}$ is electrostatic interaction energy between colloid 1 and colloid 2 at a distance $\ell$, $\varepsilon$ is the relative permittivity of the medium, $\zeta_{c1}$ is the zeta potential of colloid 1, $\zeta_{c2}$ is the zeta potential of colloid 2, $\kappa$ is the Debye-Huckel parameter.

Debye-Huckel parameter ($\kappa$ in $m^{-1}$) is determined according to equation 19 and 20 for NaCl which is a 1:1 electrolyte.

$$\kappa = \left[\frac{2000\,N_A e^2\,I}{\varepsilon_r\varepsilon_o\,kT}\right]^{1/2} \qquad 19$$

$$I = \frac{1}{2}\sum_{i=1}^{N} Z_i^2\,C_i \qquad 20$$

Where, $e$ is the elementary electric charge (1.602 x $10^{-19}$ C), $\varepsilon_r$ is relative permittivity of the electrolyte solution, $\varepsilon_o$ is the permittivity of vacuum (8.854 x $10^{-12}$ C $V^{-1}m^{-1}$), $k$ is the Boltzmann constant (1.38 x $10^{-23}$ J $K^{-1}$), $T$ is the absolute temperature (in Kelvin), $I$ is the ionic strength of the medium, $Z_i$ is the valency of the ion and $C_i$ is the molar concentration of the ion.

**Supplementary Section 3**

**Non-electrostatic Interaction Energy (NEIE) between AuNP**

To assess the nature of nanoparticle interactions due to non-electrostatic interactions, the total energy of electrodynamic interactions (NEIE, $\Delta G_{N.w}$) between the AuNP suspended in water was estimated using equation 21.[12] It should be noted here that NEIE can also be used to characterize nanoparticles in terms of their surface hydrophobicity. For hydrophobic nanoparticles values of $\Delta G_{N.w}$ < 0 and for hydrophilic nanoparticles $\Delta G_{N.w}$ > 0.

$$\Delta G_{N.w} = -2(\gamma_N^{LW} - \gamma_W^{LW})^2 - 4\left(\sqrt{\gamma_N^+ \gamma_N^-} + \sqrt{\gamma_w^+ \gamma_w^-} - \sqrt{\gamma_N^+ \gamma_w^-} - \sqrt{\gamma_w^+ \gamma_N^-}\right) \quad 21$$

NEIE of nanoparticles is an interplay of different interaction energies between nanoparticle and solvent which is water in this case. These interaction energies include:

(1). $4\sqrt{\gamma_w^+ \gamma_N^-}$ – polar adhesive energy between electron-acceptors of water molecule and electron-donors of the AuNP surface

(2). $4\sqrt{\gamma_N^+ \gamma_w^-}$ – polar adhesive energy between electron-acceptors of AuNP surface and electron-donors of the water molecules

(3). $-4\sqrt{\gamma_w^+ \gamma_w^-}$ – polar cohesive energy between electron-acceptors and electron-donors of water molecules

(4). $-4\sqrt{\gamma_N^+ \gamma_N^-}$ – polar cohesive energy between electron-acceptors and electron-donors of the AuNP surface

(5). $-2\,(\gamma_N^{LW} - \gamma_W^{LW})^2$ – apolar adhesive (Lifshitz-van der Waals) interaction energy between nanoparticles and water molecules

**NEIE between proteins and AuNP**

The nature of the non-electrostatic interaction between proteins and AuNP can be further understood by determining the NEIE between them. For two colloids suspended in water, NEIE between protein and AuNP was estimated by equation 22 where $\gamma_N^{LW}$, $\gamma_P^{LW}$ and $\gamma_w^{LW}$ are the dispersive components of surface tension; $\gamma_N^+$, $\gamma_P^+$ and $\gamma_w^+$ are the electron acceptor components of surface tension; and $\gamma_N^-$, $\gamma_P^-$ and $\gamma_w^-$ are the electron donor components of surface tension of AuNP, protein, and water respectively.

$$\Delta G_{N.w.P} = 2\left[\sqrt{(\gamma_N^{LW}\gamma_w^{LW})} + \sqrt{(\gamma_P^{LW}\gamma_w^{LW})} - \sqrt{(\gamma_N^{LW}\gamma_P^{LW})} - \gamma_w^{LW} - \left(\sqrt{\gamma_P^{LW}} - \sqrt{\gamma_w^{LW}}\right)^2 + \sqrt{\gamma_w^+}\left(\sqrt{\gamma_N^-} + \sqrt{\gamma_P^-} - \sqrt{\gamma_w^-}\right) + \sqrt{\gamma_w^-}\left(\sqrt{\gamma_N^+} + \sqrt{\gamma_P^+} - \sqrt{\gamma_w^+}\right) - \sqrt{\gamma_N^+\gamma_P^-} - \sqrt{\gamma_N^-\gamma_P^+}\right] \quad 22$$

The physical interpretation of terms in equation 22 is as follows:

(1). $-2\sqrt{\gamma_N^-\gamma_P^+}$ - Polar adhesion between electron donor component of AuNP surface tension and electron acceptor component of protein surface tension.

(2). $-2\sqrt{\gamma_N^+\gamma_P^-}$ - Polar adhesion between electron acceptor component of AuNP surface tension and electron donor component of protein surface tension.

(3). $2\sqrt{\gamma_w^-}\left(\sqrt{\gamma_N^+} + \sqrt{\gamma_P^+} - \sqrt{\gamma_w^+}\right)$ - Polar adhesion between electron acceptor groups of AuNP and proteins and electron donor ability of water molecules.

(4). $2\sqrt{\gamma_w^+}\left(\sqrt{\gamma_N^-} + \sqrt{\gamma_P^-} - \sqrt{\gamma_w^-}\right)$ - Polar adhesion between electron donor groups of AuNP and proteins and electron acceptor ability of water molecules.

(5). $2\left(\sqrt{(\gamma_N^{LW}\gamma_w^{LW})} + \sqrt{(\gamma_P^{LW}\gamma_w^{LW})} - \sqrt{(\gamma_N^{LW}\gamma_P^{LW})} - \gamma_w^{LW}\right)$ - Apolar interaction between AuNP and proteins. This can be further resolved into the following:

(I). $-2\,\gamma_w^{LW}$- apolar cohesion between water molecules

(II). $-2\sqrt{(\gamma_N^{LW}\gamma_P^{LW})}$ – apolar adhesion between AuNP and proteins

(III). $2\sqrt{(\gamma_N^{LW}\gamma_w^{LW})}$ – apolar adhesion between AuNP and water molecules

(IV). $2\sqrt{(\gamma_P^{LW}\gamma_w^{LW})}$ – apolar adhesion between protein and water molecules

### Methods

#### Synthesis of gold nanoparticles (AuNP)

Citrate reduction method[13] was modified for the synthesis of quasi-spherical citrate-capped AuNP. Briefly, 49.55 mL of trisodium citrate solution (5.4 mM) was heated at 100 °C for 10 minutes under reflux conditions. The solution was stirred using a magnetic stirrer (RCT Basic, IKA®, India). To this 150 µL of $HAuCl_4$ solution (100 mM) was added and allowed to react for 10 minutes. Following this, the temperature of oil bath was reduced to 90 °C followed by the addition of a second aliquot of 150 µL of $HAuCl_4$ solution (100 mM) to the reaction. The reaction was then allowed to proceed for 20 minutes. Finally, a third aliquot of 150 µL of $HAuCl_4$ solution (100 mM) was added to the reaction. The reaction was further allowed to proceed for another 20 minutes. After completion of the reaction, the oil bath was removed and AuNP suspension cooled for 30 minutes under mild stirring at room temperature. The final suspension of citrate-capped AuNP was transferred to a sterile polypropylene tube and stored at 4 °C until further use.

#### Characterization of AuNP

**Morphology.** The morphology of as-prepared AuNP was studied using a transmission electron microscope. For TEM samples, the nanoparticle suspension was dried on a glass slide for 2 hours at room temperature. The precipitate from the edge of the drop was collected on a Formvar-coated 300 mesh copper grid (Tedpella Inc., USA). The samples were then analyzed using a Tecnai G2 12 Twin TEM (FEI, USA) operating at 120 kV.

**Size.** The hydrodynamic diameter of AuNP was estimated by photon correlation spectroscopy using a Zetasizer ZS90 (Malvern Instruments Ltd., UK) fitted with a laser diode of 633 nm wavelength. Briefly, 1 mL sample of as prepared AuNP suspension was added to a disposable cuvette, and the sample was equilibrated at 25 °C for 180 seconds. The time-dependent decay in the intensity of light was measured and converted into a correlogram that was used to estimate the hydrodynamic diameter of AuNP using the general mode of analysis in the Zetasizer software.

**Zeta potential.** The zeta potential of as-prepared AuNP was estimated by Laser Doppler Velocimetry (LDV) using Zetasizer ZS90. LDV measures electrokinetic behavior of nanoparticles under an externally applied electric field.

**Salt-induced agglomeration of AuNP.** The stability of AuNP in increasing ionic strength of media was studied using UV-vis spectrophotometric analysis. Briefly, 50 µL of as prepared AuNP suspension was dispensed into a 96-well plate containing different concentrations of NaCl. The absorbance of AuNP suspended in different concentrations of NaCl was measured from 400-800 nm using Synergy H4 multi-mode reader at 1, 50, 100 and 150 min. The stability of AuNP was studied according to equation 23. The stability index values range from 1 (stable) to 0 (unstable).

$$Stability\ Index = \frac{\sum_{400}^{599} Absorbance_{[NaCl]} \Big/ \sum_{600}^{800} Absorbance_{[NaCl]}}{\sum_{400}^{599} Absorbance_{[control]} \Big/ \sum_{600}^{800} Absorbance_{[control]}} \qquad 23$$

**Modification and characterization of Bovine Serum Albumin (BSA)**

**Synthesis of cationic BSA (cBSA).** The surface of BSA molecules was cationized by the amine-modification of carboxyl groups of acidic amino acid residues (aspartic acid and glutamic acid).[14] Briefly, 1,6-hexanediamine was diluted in water, and the pH of the solution was adjusted to 6.5 using 1M hydrochloric acid. The resulting solution was added drop-wise to a solution of BSA under stirring conditions, and 1-Ethyl-3-(3-dimethylaminopropyl) carbodiimide (EDC) was used as a catalyst for the reaction. The reaction mixture was stirred for 5 hours at room temperature, following which pH was adjusted to 6.5 and EDC was added once again. The reaction was allowed to proceed under stirring for 6 hours. After the completion of the reaction, the solution was dialyzed against Type I water at 4 °C for 48 hours. The dialyzed cBSA solution was then stored at 4 °C.

**The secondary structure of proteins**. The secondary structure of proteins was measured using a JASCO J-815 Circular Dichroism (CD) Spectropolarimeter, fitted with a JASCO PTC-423 S/15 thermostatic cell holder for temperature control. The instrument was purged with nitrogen gas before use and a constant nitrogen flow rate of 5 liters/minute was maintained during the experiments. The protein samples were taken in a quartz cuvette of 1 mm pathlength. The temperature of the sample was adjusted to 25 °C and the ellipticity was recorded between 190 to 250 nm wavelength. The ellipticity values were converted from machine units to $\Delta\varepsilon$ by normalizing to protein concentration, mean residue weight, and pathlength of the cuvette. DichroWeb[15] was used to predict the secondary structure components of the proteins by using CONTIN algorithm[16] to fit the measured spectra to a reference dataset.

**Zeta potential.** Zeta potential of proteins was estimated in type 1 water using a Zetasizer Nano ZS90.

**Isoelectric point.** Solutions of BSA and cBSA of 10 mM ionic strength were prepared using NaCl. Protein solutions were titrated using MPT-2 Autotitrator (Malvern Instruments Ltd., UK) fitted with a degassing unit and their zeta potential was measured using a ZetaSizer Nano ZS90. The zeta potential at each pH point was determined by averaging three measurements of 40 runs each. The sample was titrated between pH 2.0 to 12.0 at an interval of 0.5 using 0.15 M NaOH, 0.15 M HCl, and 0.015 M HCl.

**Surface amine group estimation.** The relative concentration of surface amine groups of proteins was estimated by o-phtalaldehyde (OPA) assay[17] with minor modifications. Briefly, the concentration of proteins was estimated by measuring their absorbance at 280 nm using a spectrophotometer (Synergy H4 Biotek, USA). Protein sample (50 µL) was taken in a 96 well black plate (Corning, USA) and 250 µL of OPA solution (10 mg OPA, 100 µL ethanol, 5 µL of β-mercaptoethanol, and 10 mL of 0.1 M carbonate buffer of pH 10.5) was added to it. The reaction mixture was stirred at medium speed for 2 minutes and the fluorescence was measured at excitation and emission wavelengths of 340 and 455 nm respectively. The fluorescence intensity corresponded to the amine group concentration at the surface of the protein.

**Nanoparticle and protein surface characterization.**

The surface characteristic of nanoparticles and proteins was estimated by studying their partitioning behavior in three different aqueous biphasic systems (ABS). The theoretical framework developed to correlate partitioning coefficient of nanoparticles, which is an experimentally observable quantity, to the surface characteristics of nanoparticles is provided in **Supplementary Section 1**. Three ABS listed in **Table S5** were optimized to achieve equal volumes of both the top and bottom phases. The ABS reported here were prepared from a single batch of polymers. As natural polymers show a batch-to-batch variation, we recommend that ABS should be optimized to obtain equal volume every time a new batch of polymer is procured. Also, the ABS are temperature sensitive and utmost care should be taken to minimize the temperature variations.

**Table S5**. Aqueous biphasic systems (ABS) used for the partitioning of colloids. ABS were prepared at 25 °C.

| | **Polymer 1** | **Volume (µL)** | **Polymer 2** | **Volume (µL)** | **Water (µL)** | **AuNP* (µL)** |
|---|---|---|---|---|---|---|
| ABS1 | PEG 1000 (40% w/w) | 340 | Dextran 40000 (40% w/w) | 400 | 210 | 50 |
| ABS 2 | PEG 6000 (40% w/w) | 160 | Dextran 40000 (40% w/w) | 300 | 490 | 50 |
| ABS 3 | PEG 10000 (40% w/w) | 145 | Dextran 40000 40% w/w) | 300 | 505 | 50 |
| *concentration = 180 µg $mL^{-1}$ | | | | | | |

**Partitioning of AuNP.** For studying the partitioning behavior of AuNP in ABS, the desired volume of PEG and dextran solutions were added to a 2 mL micro-centrifuge tube. To this 50 µL of as prepared AuNP suspension was added, and the final volume was made up to 1000 µL using type 1 water (resistivity 18 MΩ cm). The contents of micro-centrifuge tubes were then mixed thoroughly. ABS were then incubated for 24 hours at 25 °C. After 24 hours, 200 µL of samples were drawn from the top phase of ABS, and the amount of AuNP present in that phase was estimated by measuring absorbance at 450 nm in a 96 well plate using Synergy H4 multimode reader (Biotek, USA). The wavelength of 450 nm was chosen as the surface plasmon resonance peak of AuNP is sensitive to the polarity of the surrounding medium. The optical density was corrected to 1 cm pathlength by using equation 24. The pathlength correction factor ($l$) was calculated as the ratio of the difference in the optical density at 900 and 977 nm of the sample to that of blank phases measured in a 1 cm reference cell in Take 3® plates (equation 25).

$$A_{1cm} = \frac{A_{200\mu L}}{l} \tag{24}$$

$$l = \frac{(A_{977} - A_{900})_{sample}}{(A_{977} - A_{900})_{reference,1.0\ cm}} \tag{25}$$

The partition coefficient (Equation26) and $\Delta\gamma$ (Equation 27) were then calculated from the corrected optical density values. For ease of calculation, it was assumed that the particle partition had taken place between top and bottom phases as the interfacial tension between the phases is very small and therefore will not have a significant contribution in estimating $\Delta\gamma$ value.

$$K_{TB} = \frac{A_{450,TP}}{A_{450,water} - A_{450,TP}} \tag{26}$$

$$\ln K_{TB} = -\frac{\pi\, D_{AuNP}^2\, \Delta\gamma}{kT} = -\frac{A\, \Delta\gamma}{kT} \tag{27}$$

**Partitioning of proteins.** Protein partitioning coefficient was estimated by measuring the protein concentration in top and bottom phase of ABS using o-phthalaldehyde (OPA) assay. Briefly, 100 µL of protein solution was added to the ABS described in **Table S5** while maintaining the volume at 1000 µL. After partitioning a 50 µL sample was carefully retrieved from the top and bottom phases of ABS and added to a 96-well plate. To this, 250 µL of OPA assay solution was added and the plate was stirred for 2 min at a medium speed in a multimode spectrophotometer (Synergy H4, Biotek, USA). The fluorescence was measured at excitation and emission wavelengths of 340 and 455 nm respectively. The partition coefficient of proteins was measured as a ratio of fluorescence intensities measured for the top and bottom phases (equation 28).

$$K_{TB} = \frac{I_{top\ phase}}{I_{bottom\ phase}} \tag{28}$$

**Experimental framework for the resolution of the surface tension of polymeric phases**

The surface tension of a liquid or a condensed matter is further resolved into dispersive/Lifshitz-van der Waals ($\gamma_L^{LW}$) and polar/acid-base ($\gamma_L^{AB}$). $\gamma_L^{LW}$ interactions are van der Waals forces which include non-covalent, non-electrostatic, apolar, electrohydrodynamic intermolecular interactions such as Keesom energies, Debye energies, and London forces. The rate of decay of all of these interfacial interactions as a function of inter-particle distance is same at a macroscopic level.[18] $\gamma_L^{AB}$ is resolved further into the electron acceptor/acidic ($\gamma^+$) and the electron donor/basic ($\gamma^-$) parameters (equation 29).

$$\gamma_L = \gamma_L^{LW} + \gamma_L^{AB} = \gamma_L^{LW} + 2\sqrt{\gamma_L^+\gamma_L^-} \qquad 29$$

The surface tension of polymeric phases was resolved into various components as per the experimental framework provided below.

**Surface tension analysis.** The surface tension of top ($\gamma_T$) and bottom ($\gamma_B$) phases was measured using pendant drop method. Each phase of ABS was carefully separated while keeping the interfacial layer unperturbed. The polymeric solutions were then used to measure surface tension with the help of a goniometer (OCA-35, DataPhysics, Germany). The solution was pumped with a metered pump through a blunt stainless-steel needle of 1.6 mm diameter till a drop was about to fall under the influence of gravity. Large needle size was used in the determination of pendant drop as it causes less variation in the estimated value of surface tension.[19] The drop profile was then analyzed using the software provided with the goniometer to determine the surface tension of the phases of ABS.

**Resolution of surface tension into polar and dispersive components.** Surface tension was further resolved into $\gamma_L^{LW}$ and $\gamma_L^{AB}$ components of surface tension by measuring the contact angle of polymeric phases on polyethylene thin films. Polyethylene is an apolar polymer (dispersive component of surface energy of 35.7 mJ m$^{-2}$ at 20 °C).[20] Low-density polyethylene (LDPE) of Mn 7,000 (Sigma-Aldrich, USA) was used for thin film preparation.

1) **LDPE thin film preparation.** LDPE thin films were prepared by heat press method to avoid using solvent mediated artifacts while measuring contact angle. Briefly, rectangular

glass coverslips were cleaned thoroughly using Piranha Solution (3 parts concentrated sulphuric acid and 7 parts of concentrated hydrogen peroxide) and then washed with copious amount of type 1 water in an ultrasonic water bath (Equitron, MIMC, India). These coverslips were dried in a hot air oven in a clean glass petri dish. To prepare LDPE thin films, a coverslip was heated to a temperature of 100 °C, and a small amount of polymer was placed on heated coverslip followed by placing a clean coverslip on the top of polymer. The polymer was melted in between the coverslips to form a uniform thin film. LDPE thin films were stored under a nitrogen atmosphere until further use.

2) **Contact angle measurement of phases of ABS.** Contact angles of phases of ABS were measured on LDPE thin film using a goniometer (OCA-35, DataPhysics, Germany). Briefly, Hamilton repeating dispenser fitted with 250 µL Gastight syringe (Hamilton Robotics, USA) was used to dispense 5 µL drop of the phase of ABS onto the LDPE thin films which were cleaned with a sterile nitrogen jet prior to use. These drops were equilibrated for a minute, and then the goniometer was used to image the drop. The image profile was fitted with the help of an in-built software to estimate the contact angle of the phase on LDPE. The contact angle was used to estimate the $\gamma_L^{LW}$ component of the surface tension using equation 30.

$$\gamma_L(1+\cos\theta) = 2\sqrt{\gamma_L^{LW}\gamma_S^{LW}} \qquad 30$$

**Resolution of $\gamma_L^{AB}$ into $\gamma_L^-$ and $\gamma_L^+$**

A squared relationship between surface tension components and Kamlet-Taft parameters has been implicated in literature (Equation 31).[21-22] Therefore, the squared relationship between Kamlet-Taft's acidity/basicity value and electron donor ($\gamma_L^-$)/acceptor ($\gamma_L^+$) components of surface tension could be used to resolve polar component of surface tension into $\gamma_L^-$ and $\gamma_L^+$ components as per equation 32.

$$x = \frac{\gamma^+}{\gamma^-} = \left(\frac{\alpha}{\beta}\right)^2 \qquad 31$$

$$\gamma_L^{AB} = 2\,\gamma_L^-\sqrt{x} = 2\,\gamma_L^-\left(\frac{\alpha}{\beta}\right) \qquad 32$$

**Solvatochromic analysis of phases.** Solvent properties of polymeric phases of ABS were analyzed using solvatochromic analysis.[23] Solvatochromic analysis yields information about the following properties of a solvent (Kamlet-Taft parameters)[24]

1. **Polarity/polarizability parameter (π*):** $\pi^*$ is a measure of dipole-dipole and dipole-induced dipole interactions among solute and solvent molecules.[24] For determining polarity/polarizability parameter ($\pi^*$), spectroscopic shift in absorption maxima of 4-nitroanisole was used. The wavenumber of absorption maxima ($\nu_{max}$) was determined in units of 1000 $cm^{-1}$ and its value was used in equation 33.[25-26] For calculation of $\pi^*$, $\nu_{cyclohexane}$ was calculated by assuming $\pi^*_{cyclohexane} = 0$ and $\nu_{DMSO}$ was calculated by assuming $\pi^*_{DMSO} = 1$. The absorption spectrum was measured from 250-400 nm.

$$\pi^*(solvent) = \frac{\nu_{max,NA} - \nu_{cyclohexane}}{\nu_{DMSO} - \nu_{cyclohexane}} = \frac{\nu_{max,NA} - 34.12}{31.72 - 34.12} \qquad 33$$

2. **Hydrogen bond acceptance (HBA)/ electron pair donation ability to form a coordinative bond (β):** For determining HBA (β), a pair of 4-nitrophenol and 4-nitroanisole was used. HBA of solvent was calculated from the wavenumber (in units of 1000 $cm^{-1}$) of absorption maxima of 4-nitroanisole and 4-nitrophenol as per equation 34.[27]

$$\beta = \frac{-\Delta\Delta\nu_{(NP,NA)}}{2.31} = \frac{\left(0.901\,\nu_{max,NA} + 4.16 - \nu_{max,NP}\right)}{2.31} \qquad 34$$

3. **Hydrogen bond donation (HBD) ability (α):** Equation 35 was used for estimating the HBD of polymeric phases.[28] Betaine is insoluble in water, therefore, a water soluble dichloro-substituted betaine dye [2,6-dichloro-4-(2,4,6-triphenyl-N-pyridinio)-phenolate] was used for estimation of $\nu_{max,Betaine}$. For this, wavenumber (in units of 1000 $cm^{-1}$) of dichlorobetaine was measured in polymeric phases and this wavenumber was used for estimating $\nu_{max,Betaine}$ using equation 36. Equation 36 was obtained by exponentially fitting the data of wavenumber of absorption maxima of betaine and dichlorobetaine for different solvent systems as reported by Kessler and Wolfbeis.[29]

$$\alpha = \frac{\nu_{max,Betaine} + 1.873\ \nu_{max,NA} - 75.58}{6.25} \quad 35$$

$$\nu_{max,Betaine} = 6.75 + 1.33\ e^{0.1\ \nu_{max,Dichlorobetaine}} \quad 36$$

The important point is to use same batch of polymers to obtain the resolved components of surface tension and to perform ABS partitioning of nanoparticles and biomacromolecules. This is to avoid erroneous results due to batch-to-batch variation of natural polymers procured from same commercial source.

**Interaction energy calculations.**

Non-electrostatic interaction energies (NEIE) between AuNP-AuNP and AuNP-protein was calculated using a set of equations described in **Supplementary Section 3**. The inter-particle distance dependent decay in the total interaction energy between AuNP or AuNP-protein was calculated using extended DLVO (xDLVO) theory as outlined in **Supplementary Section 2**.

**Effect of nanoparticle-protein interaction on secondary structure of proteins**

The effect of nanoparticle-protein interaction on the secondary structure of the proteins was studied using CD spectroscopy.[30] Briefly, 500 µL of as prepared AuNP was incubated with 500 µL of 0.4 mg $mL^{-1}$ solution of BSA or cBSA for 24 hours at 25 °C. The ellipticity of proteins in presence and absence of AuNP was measured using a Circular Dichroism spectropolarimeter (JASCO J-815), fitted with a thermostatic cell holder (JASCO PTC-423 S/15). The temperature of the sample was adjusted from 25 °C to 80 °C, with an interval of 5 °C, and ellipticity between 190 to 250 nm wavelength was measured at each temperature point. Data from three such measurements were averaged for each sample. The CD data was then used to calculate thermodynamics of unfolding of the proteins. First, the fraction of protein unfolded, $K_{eq}$ at a given temperature, $T$, was calculated using equation 37 where $\theta_T$ is the ellipticity of protein at a given temperature, $\theta_F$ is the ellipticity of fully folded protein, and $\theta_U$ is the ellipticity of protein in fully unfolded state. For calculations ellipticity of native proteins at 25 °C was used as $\theta_F$ and ellipticity of native and AuNP-adsorbed proteins at 80 °C was considered as $\theta_U$. The enthalpy and entropy of protein unfolding in presence and absence of AuNP was estimated using van't Hoff equation (equation 38). The melting temperature, $T_m$, is the temperature at which the free energy of protein unfolding is zero and was estimated according to equation 39.

$$K_{eq} = \frac{[U]}{[F] + [U]} = \frac{\theta_F - \theta_T}{\theta_F - \theta_U} \quad 37$$

$$\ln K_{eq} = -\frac{\Delta H}{RT} + \frac{\Delta S}{R} \quad 38$$

$$T_m = \frac{\Delta H}{\Delta S} \quad 39$$

### Supplementary References


1. van Oss, C. J. Lifshitz–van der Waals and Lewis Acid-Base Properties of Liquid Water - Physical and Physico-Chemical Effects. In *Interfacial Forces in Aqueous Media, Second Edition*; Taylor & Francis; 2006. Pp 93-99.

2. Abraham, M. H. Scales of Solute Hydrogen-Bonding: Their Construction and Application to Physicochemical and Biochemical Processes. *Chem. Soc. Rev.* **1993,** *22*, 73-83.

3. Luechau, F.; Ling, T. C.; Lyddiatt, A. A Descriptive Model and Methods for Up-Scaled Process Routes for Interfacial Partition of Bioparticles in Aqueous Two-Phase Systems. *Biochem. Eng. J.* **2010,** *50*, 122-130.

4. Albertsson, P. A. The Distribution of Particles in a Two-Phase System - Theory. In *Partition of Cell Particles and Macromolecules: Distribution and Fractionation of Cells, Viruses, Microsomes, Proteins, Nucleic Acids, and Antigen-Antibody Complexes in Aqueous Polymer Two-Phase Systems*; J. Wiley; 1960; pp 95-107.

5. Schrader, M. E. Young-Dupre Revisited. *Langmuir* **1995,** *11*, 3585-3589.

6. Hiemenz, P. C.; Rajagopalan, R. Surface Tension and Contact Angle: Application to Pure Substances. In *Principles of Colloid and Surface Chemistry, Third Edition, Revised and Expanded*; Taylor & Francis; 1997; pp 248-296.

7. van Oss, C. J.; Ju, L.; Chaudhury, M. K.; Good, R. J. Estimation of the Polar Parameters of the Surface Tension of Liquids by Contact Angle Measurements on Gels. *J. Colloid Interface Sci.* **1989,** *128*, 313-319.

8. Brant, J. A.; Childress, A. E. Membrane-Colloid Interactions: Comparison of Extended DLVO Predictions with AFM Force Measurements. *Environ. Eng. Sci.* **2002,** *19*, 413-427.

9. van Oss, C. J. Theory. In *Interfacial Forces in Aqueous Media, Second Edition*; Taylor & Francis: 2006; pp 9-89.

10. Bos, R.; Van Der Mei, H. C.; Busscher, H. J. Physico-Chemistry of Initial Microbial Adhesive Interactions - Its Mechanisms and Methods for Study. *FEMS Microbiol. Rev.* **1999,** *23*, 179-229.

11. Sader, J. E.; Carnie, S. L.; Chan, D. Y. Accurate Analytic Formulas for the Double-Layer Interaction Between Spheres. *J. Colloid Interface Sci.* **1995,** *171*, 46-54.

12. van Oss, C. J. Hydrophobicity of Biosurfaces - Origin, Quantitative Determination and Interaction Energies. *Colloids Surf., B* **1995,** *5*, 91-110.

13. Frens, G. Controlled Nucleation for the Regulation of the Particle Size in Monodisperse Gold Suspensions. *Nature* **1973,** *241*, 20-22.

14. Huang, X.; Li, M.; Green, D. C.; Williams, D. S.; Patil, A. J.; Mann, S. Interfacial Assembly of Protein–Polymer Nano-Conjugates into Stimulus-Responsive Biomimetic Protocells. *Nat. Commun.* **2013,** *4*, ncomms3239.

15. Whitmore, L.; Wallace, B. A. Protein Secondary Structure Analyses from Circular Dichroism Spectroscopy: Methods and Reference Databases. *Biopolymers* **2008,** *89*, 392-400.

16. Provencher, S. W.; Gloeckner, J. Estimation of Globular Protein Secondary Structure from Circular Dichroism. *Biochemistry* **1981,** *20*, 33-37.

17. Benson, J. R.; Hare, P. O-Phthalaldehyde: Fluorogenic Detection of Primary Amines in the Picomole Range. Comparison with Fluorescamine and Ninhydrin. *Proc. Natl. Acad. Sci. U. S. A.* **1975,** *72*, 619-622.

18. van Oss, C. J. Lifshitz–van der Waals (LW) Interactions. In *Interfacial Forces in Aqueous Media, Second Edition*; Taylor & Francis; 2006. pp 9-18.

19. Lin, S.-Y.; Wang, W.-J.; Lin, L.-W.; Chen, L.-J. Systematic Effects of Bubble Volume on the Surface Tension Measured by Pendant Bubble Profiles. *Colloids Surf., A* **1996,** *114*, 31-39.

20. Wu, S. Surface and Interfacial Tensions of Polymer Melts: I. Polyethylene, Polyisobutylene, and Polyvinyl Acetate. *J. Colloid Interface Sci.* **1969,** *31*, 153-161.

21. Della Volpe, C.; Siboni, S. Some Reflections on Acid–Base Solid Surface Free Energy Theories. *J. Colloid Interface Sci.* **1997,** *195*, 121-136.

22. Della Volpe, C.; Siboni, S. Acid–Base Surface Free Energies of Solids and the Definition of Scales in the Good–van Oss–Chaudhury Theory. *J. Adhes. Sci. Technol.* **2000,** *14*, 235-272.

23. Marcus, Y. The Properties of Organic Liquids That are Relevant to Their Use as Solvating Solvents. *Chem. Soc. Rev.* **1993,** *22*, 409-416.

24. Heger, D.; Klán, P. Interactions of Organic Molecules at Grain Boundaries in Ice: A Solvatochromic Analysis. *Journal of Photochemistry and Photobiology A: Chemistry* **2007,** *187*, 275-284.

25. Kamlet, M. J.; Abboud, J. L.; Taft, R. W. The Solvatochromic Comparison Method. 6. The π Scale of Solvent Polarities. *J. Am. Chem. Soc.* **1977,** *99*, 6027-6038.

26. Laurence, C.; Nicolet, P.; Dalati, M. T.; Abboud, J.-L. M.; Notario, R. The Empirical Treatment of Solvent-Solute Interactions: 15 Years of π*. *The Journal of Physical Chemistry* **1994,** *98*, 5807-5816.

27. Kamlet, M. J.; Taft, R. W. The Solvatochromic Comparison Method. I. The β-Scale of Solvent Hydrogen-Bond Acceptor (HBA) Basicities. *J. Am. Chem. Soc.* **1976,** *98*, 377-383.

28. Taft, R. W.; Kamlet, M. J. The Solvatochromic Comparison Method. 2. The α-Scale of Solvent Hydrogen-Bond Donor (HBD) Acidities. *J. Am. Chem. Soc.* **1976,** *98*, 2886-2894.

29. Kessler, M. A.; Wolfbeis, O. S. ET(33), A Solvatochromic Polarity and Micellar Probe for Neutral Aqueous Solutions. *Chem. Phys. Lipids* **1989,** *50*, 51-56.

30. Greenfield, N. J. Using Circular Dichroism Collected as a Function of Temperature to Determine the Thermodynamics of Protein Unfolding and Binding Interactions. *Nat. Protoc.* **2006,** *1*, 2527-2535.
